# Improving the benchmark of variant calling in clonal bacteria using more realistic *in silico* genomes, the case of *Mycobacterium tuberculosis*

**DOI:** 10.1101/2025.10.04.680280

**Authors:** Adrien Le Meur, Ricardo C. Rodriguez de la Vega, Rima Zein-Eddine, Guislaine Refrégier

## Abstract

The democratisation of Whole Genome Sequencing data in bacterial genomics requires the benchmarking of associated analytical methodologies such as reference-based variant calling. Current variant calling benchmarks rely either on *de novo* assembled natural genomes, for which true variants are inferred using a genome aligner, or on genomes evolved *in silico* by incorporating short variants on reference genomes. We introduce Maketube, a method for evolving realistic genomes of the *Mycobacterium tuberculosis* complex with the full diversity of variants verified in natural isolates, and describe benchmarking results using Maketube-evolved genomes.

We document that Maketube-evolved genomes satisfyingly mimic Mtbc complex genomes. Using Maketube-evolved genomes, we show that genome aligners miss up to 7.5% of the variants, which implies that benchmarkings with natural *de novo* assembled genomes are biased. Second, we show that recall of popular variant calling pipelines MTBseq, TB-Profiler, and our in-house genomic pipeline genotube, was overestimated by 1 to 10% in benchmarkings relying on simplistic *in silico-*evolved genomes, and that slight but significant differences in performance exist between pipelines. Finally, we provide evidence that variants are missed in duplicated regions and in regions flanking sequences absent in the reference (displaced insertion sequences or sequences deleted during the evolution of the reference).

Altogether, realistic *in silico*-evolved genomes such as Maketube-derived ones are precious tools for reliable genomic tools benchmarking. We provide new evidence that structural variants interfere with variant-calling, both because of the additional sequences they contain, but also because of misalignments around insertions.

## Background

Although long-read whole-genome sequencing (WGS) is increasingly reshaping the field of genomics, the majority of publicly available genomic data, particularly for relatively simple genomes such as bacterial pathogens, still originates from short-read sequencing. Thousands of samples are regularly analysed by single labs, and large databases have been set up [1]. For instance, a “*Mycobacterium tuberculosis”* query retrieved 275,865 experiment entries in in the European Nucleotide Archive (ENA) for short-read Illumina technology (reversible terminators based chemistry) and 3,501 for long-read Oxford Nanopore Technologies MinION on Oct 1^st^, 2025. New methodologies arise to extract useful information out of these data based on Artificial Intelligence (AI) [2, 3] and/or on k-mer reference-free approaches [4], that need to be compared to more standard reference-based variant calling.

Reference-based variant calling of short-read data consists in aligning reads to a reference genome, identifying where the reads locally differ from the reference (*i.e.,* identifying short variants), and filtering them to exclude likely false positives. As most phenotypes of interest, such as antibiotic resistance, are mainly controlled by short variants, this methodology efficiently summarizes the most relevant information in a new genome. The resulting information is valuable not only for research but also for clinical applications, as most antibiotic resistance conferring mutations can be reliably identified. Still, reducing sequencing data to a mere list of short variants occurs at some cost, which should be born in mind to have a fine understanding of actual genome composition [5]. Assessing the impact of the tools used for variant calling in bacterial genomes, the aim of corresponding benchmarking methodologies, has accordingly concentrated large efforts in the last 5 years [6–8].

The causative agents of human and animal typical tuberculosis belong to highly related monophyletic and paraphyletic species, named after their main host, grouped in the monophyletic taxon called the *Mycobacterium tuberculosis* complex (Mtbc), with main lineages sharing more than 99.78% Average Nucleotide Identity (ANI), and around 99.3% with *M. canettii* strains [9]. A particularly high number of publicly available samples have been sequenced since the commercialization of short-read sequencers around 2005, so that researchers from all around the world have unrestricted access to the short-read sequencing data of thousands of Mtbc strains isolated from patients all over the world [10]. The corresponding genomes differ from each other by short variants (Single Nucleotide Polymorphisms, referred to as SNPs, and small insertions and deletion, referred to as indels), and large variants also referred to as structural variants, that include regions of deletions (RDs), number and positions of insertion sequences (IS) and duplications [11]. Using different types of variants, these data have enabled the discovery of large-scale phenomena concerning genome evolution, such as variations in selection pressures through time, retrospective studies highlighting potential analytical biases, and an overall better understanding of the species diversity [12–16].

Reference-based variant calling of Mtbc, as proposed by several all-in-one pipelines, follows three standard steps: 1) mapping, where the reads are aligned on a reference genome (often the sequence of H37Rv, sequenced in 1998 and continuously polished for 15 years); 2) variant-calling *per se*, where the resulting alignment is parsed by tools such as Freebayes, GATK HaplotypeCaller, Deepvariant, and samtools’ mpileup [10, 17–20], to spot short variants relative to the reference genome; 3) filtering, where additional filters are applied to remove variants of low reliability. The all-in-one bioinformatics pipelines, whose interoperability is secured by workflow managers or centralised analysis, include: Phyresse, MTBseq, TB-Profiler, UVP and XBS [18, 19, 21–23]. They all include the prediction of antibiotic resistance and lineage identification and exhibit some specific aims: identifying clusters and inferring transmission (MTBseq, UVP, XBS) and/or handling complex samples exhibiting contamination (XBS). In addition to specific focuses, these pipelines use different variant-calling tools and include different filtering processes, with different balances between precision/sensitivity and specificity/recall. As for other pathogens, benchmarking activities have helped to identify best practices, to evaluate global performance of the tools, and have concentrated large efforts recently [7, 22, 24].

Benchmarking consists in measuring the advantages and limitations of different tools used in a process, by performing many tests and communicating their results. If benchmarking principles are straightforward, the underlying methodology is not so simple in Mtbc as for other pathogens. The first difficulty in providing a reliable benchmark of variant calling for Mtbc with data validated by the community is the availability of both reliable assembled genomes and corresponding short-read raw data. While many high-quality complete genomes are now produced thanks to the large improvements of raw data acquisition and analytic pipelines, it is dangerous to use any randomly picked data. The only community-accepted gold standards are the highly polished genomic data resulting from Sanger-sequencing, in combination with long-read and short-read sequencing, as could be performed for H37Rv and *M. bovis* AF2122-97 genomes, thanks to more than 20 years of cross-checking. This triggers a second difficulty: it is impossible to reflect Mtbc with few highly reliable/gold standard genomes alone. Two approaches were developed to incorporate other highly reliable assembled genomes into benchmarking studies. These approaches differ in the way they generate or identify the so-called true variants that are later compared with the variants detected by analysis pipelines.

The first approach for benchmarking WGS analyses is to genrenate *in silico* genomes by artificially introducing short variants in the reference genome [17, 22, 25]. Genomes are evolved using tools such as SNP-Mutator or Mutation-Simulator [26, 27], introducing artificial SNPs and indels that match the frequency and nature found in real samples. The set of true SNPs and indels is provided as a separate output by the tool. Artificial reads are then generated from the evolved genome with tools such as wgsim or ART_Illumina [28, 29]. These tools mimic most properties of real sequencing experiments, such as random errors and variable Phred scores for each read, and the error parameters can be adjusted to best correspond to real data. The artificial reads archives are processed by the pipelines to benchmark, and the detected variants are compared to the true set of variants. The limitation of this approach is that the evolved genomes perfectly align to the reference genome, *i.e.* variations other than SNPs and indels are not created, and their impact on alignment is therefore not evaluated.

The second approach to benchmarking WGS analyses’ relies on high-quality *de novo* reconstructed genomes, resulting from the combination of long-read and short-read sequencing data [6, 30]. Long reads are first assembled and polished using tools such as Flye or Circlator [31, 32], then filtered high-quality short reads are used for further polishing with for instance Pilon [33]. This methodology has, for instance, served to establish a set of true variants for Human Genomics in the “Genome In a Bottle” project [34]. The true short variants are retrieved thanks to the alignment of the reconstructed genome and reference genome using genome aligners such as MUMmer or minimap2 [35, 36]. The variants obtained with the pipelines to benchmark are compared to the true variants, as in the first approach. The limitation of this second approach is that it relies on the ability of genome aligners to identify all true variants: false positives and false negatives of these genome aligners are not taken into account which implies that the true variants might not all be spotted.

Altogether, benchmarking requires knowing reliable true variants on genomes representative of the diversity of data to analyse, here genomes of Mtbc strains. Genomes analysed in the current benchmarking have limitations in the representativity and/or in the exhaustivity of true variants. Artificial genomes more realistically representing real genomes could offer more reliable benchmarks. Methods such as Artificial Life Framework (ALF) have been developed to evolve genomes *in silico* with a higher complexity than mere short variant incorporation, using varying evolutionary models, incorporating gene losses and gene gains mimicking Lateral Gene Transfer, selection pressures, and integrating population genomics. They allow retrieval of ancestral genomes, evolutionary events specific to each branch in the evolutionary tree [37, 38], and recent models incorporate the possibility to evolve in parallel genomes with different selection pressures [39]. They allow disentangling the impact on the fixation of mutations of interest of various parameters such as mutation rate dynamics, effective population size, and selection pressure. Yet, these tools do not retain full information on the correspondence between ancestral and evolved versions of the genome in a user-friendly format.

The first objective of our study was to generate artificial Mtbc genomes evolved *in silico* using realistic evolutionary parameters and keep exhaustive track of the variant positions. We employed a methodology similar to ALF, with customized settings designed to closely replicate the evolutionary dynamics of real genomes. This approach includes a detailed characterization of evolutionary features and allows for precise backtracking to ancestral genome positions. The main purpose of the resulting genomes was to complete high-sensitivity benchmarking of variant calling pipelines classically used for Mtbc. This study was made possible thanks to the large knowledge acquired in the past few years on all types of structural variants in Mtbc genomes, including insertion sequence IS*6110* dynamics [40], regions of deletion diversity [15], and duplication characteristics [41]. Generating these new artificial genomes was performed using Maketube, an R package that inserts structural variants and short variants into a reference genome. Maketube-evolved genomes are more representative of Mtbc diversity than standard *in silico*-evolved genomes, and they do not need genome aligners to identify true variants, in contrast to long-read based *de novo* reconstructed genomes. Using Maketube genomes, we first document that genome aligners used in benchmarking based on *de novo* reconstructed genomes introduce false negatives. Second, we provide a first highly-sensitive benchmark of pipelines commonly used by the Mtbc genomics community, showing the higher precision but lower recall of MTBseq and Genotube as compared to TBprofiler, and we document the lower recall of all pipelines near regions absent in the reference.

### Data description

#### Genome sets’ description

This study includes three different sets of Mtbc genomes: sets A, B and C. Set A consists of 24 previously assembled genomes corresponding to real strains (**Fig. 1A, Suppl. data S1**), selected as high-quality natural genomes, from 5 different lineages. Specifically, set A includes five reference strains sequenced using the Sanger method (L4 CDC1551, *M. bovis* BCG Pasteur from animal lineage, L2 W-148, L4 Erdman and L4 F11), eighteen genomes built with long read sequencing and polished with short read sequencing, all with a “complete genome” status on NCBI [12 L4 assemblies from [30], 3 L2 *Beijing* from [42] and 3 unpublished L1 genomes) and one L5 from Benin [43] assembled exclusively with long read PacBio SMRT sequencing. Set B includes 180 artificial genomes, ninety evolved by SNP-Mutator [30 from L4 H37Rv, 30 from L2 Beijing 18b, 30 from bovis AF2122/97 belonging to animal lineage, a lineage p) and ninety artificial genomes evolved by Maketube (3x10=30, *i.e.* 3 populations of 10 genomes evolved from H37Rv, 3x10=30 from Beijing 18b, and 3x10=30 from bovis AF2122/97) as detailed below in the Methods (**Fig. 1B**). Set C is an independent set of 200 Maketube genomes (20×10), evolved from H37Rv, including more variants to study the impact of structural variants on reference-based variant calling (**Fig. 1C**).

**Figure 1.**
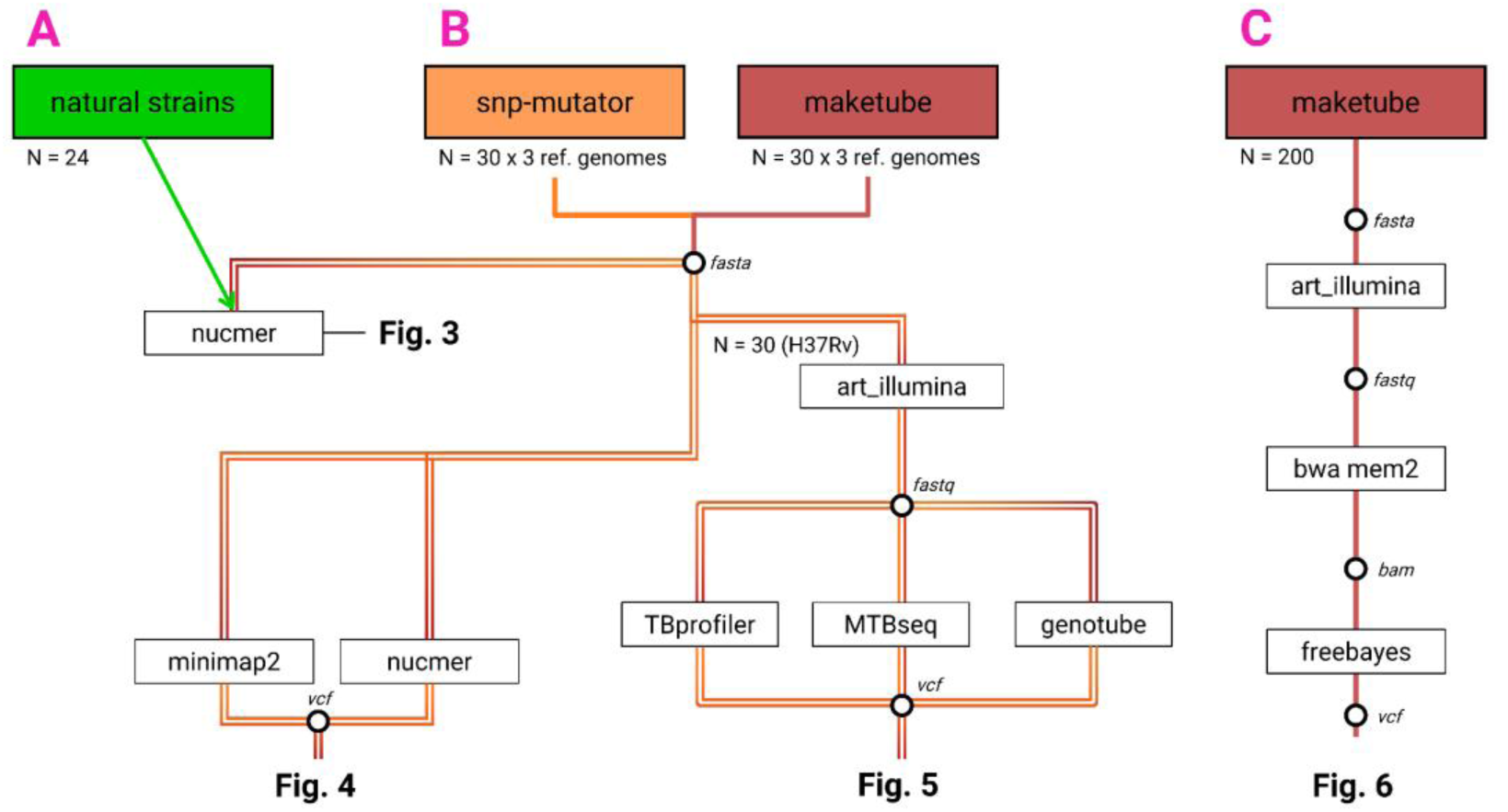
Project outline: genome sets A, B and C, and the subsequent analyses. The colored squares at the top describe the origin of the full-length genomes (reconstructed or evolved by *in silico* tools) used in the study. The numbers below indicate the total number of genomes. These genomes are fully described in **S1 table** and available on GitHub. Of note, all Maketube-evolved genomes were built by sets of 10 genomes carrying the same structural variants. Empty boxes show the implemented tools (software, pipelines). Data format is shown in italics. The figures presenting the output of the analyses are shown in bold.

## Results

### Building *in silico* genomes of Mtbc with Maketube

We developed Maketube, an R tool that evolves genomes from a single Mtbc source genome, that will also serve as a reference when performing reference-based variant calling. Maketube implements all known evolutionary events observed in real Mtbc genomes while keeping the positional correspondence of each nucleotide in the source and in the evolved genomes. Implemented evolutionary processes belong to two classes: structural variations and short variations. Structural variations include: insertion sequences movement [40], large deletions [15, 44], and duplications [41, 45]. We add a fourth structural variation to mimic the presence, in all real genomes, of accessory genome parts present in the Mtbc ancestor but deleted in the genome used for reference-based variant calling. We chose to simulate the sequences to be added by compiling kmers from real genomes. To do so, we selected reads from various strains that did not align to the reference genome, cut them into 31-mers and picked randomly among these kmers. We call these inserts “Ancestral-like regions” (**Fig. 2**, structural variant type 3, in purple). The number and sizes of the deletions and insertions were sourced from literature, *i.e.* number and/or sizes were drawn from distributions matching real data. The position and size of the duplication having no impact on the problem they specifically pose during read alignment, which is the random alignment with one of the two duplicates, we implemented a single duplication half the size of the longest described. To keep track of nucleotide positions, the types of structural variants are added successively to a reference genome (see Methods for details). Short variants are then introduced in each structurally evolved genome (∼600 SNPs as in standard sets of clinical strains in Set B, ∼2,500 SNPs in Set C). Different short variants were introduced in the same structurally-evolved genome, to mimic genomes belonging to the same population. All resulting Maketube genomes are linear genomes featuring: a displacement of all IS*6110* insertions causing no change in genome length (feature 1, green in **Fig.2**), three deletion regions as compared to H37Rv for a total length in the range of 4,011-29,156 bp (feature 2, blue in **Fig.2**, only one shown), the insertion of Ancestral-like regions to reach a total length of 0.2 to 0.8% of total genome (feature 3, purple in **Fig. 2**), and a region of duplication of 150,000 bp, with the duplicata placed at the end of the genome (feature 4, light pink in **Fig. 2**, origin of the duplicata not shown for the sake of clarity).

**Figure 2.**
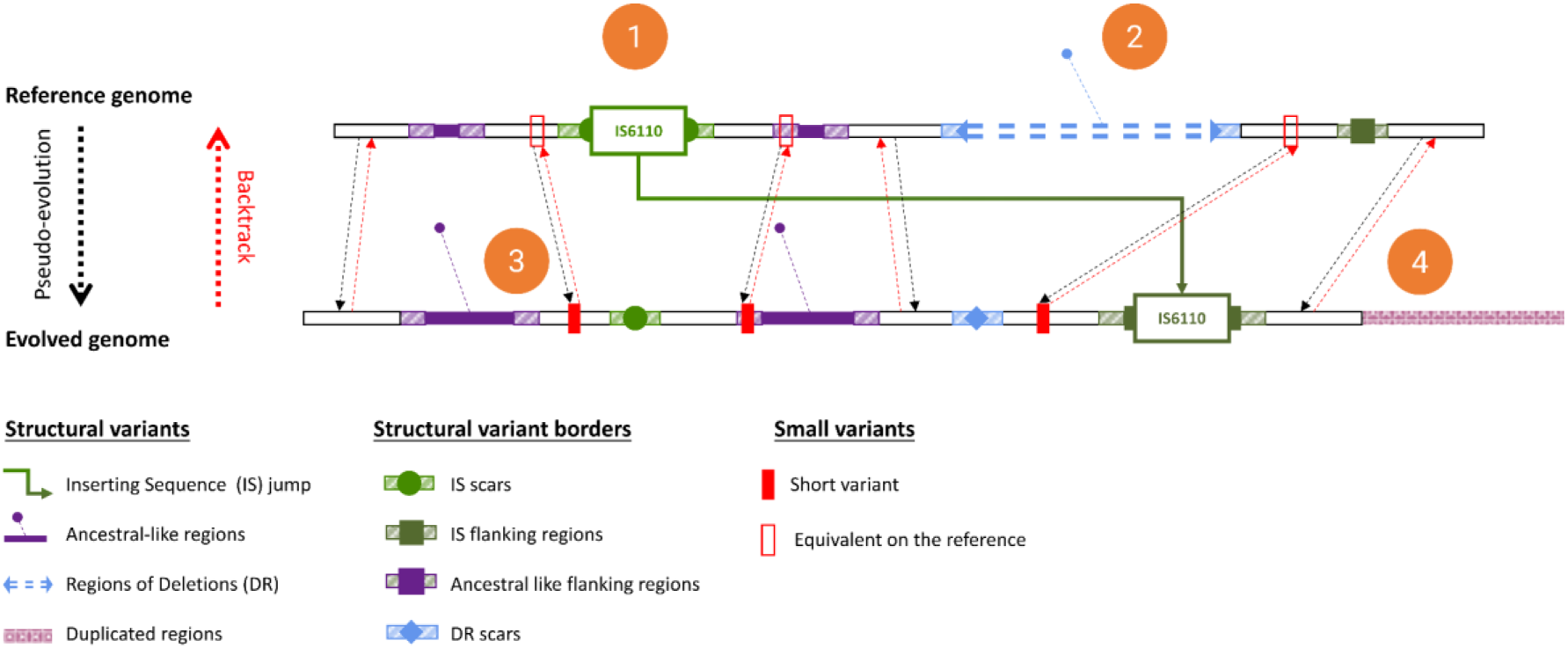
Genome evolution principles implemented in Maketube. Four types of structural variants are implemented for each simulation, numbered 1 to 4 (orange). Short variants inserted in evolved genomes are shown as solid red rectangles, with their corresponding positions in the reference indicated by empty rectangles. Correspondences between reference and evolved genomes are illustrated with black and red arrows. Borders of structural variations, which might have a higher prevalence of missed SNPs, are hatched. For details on the number and sizes of the variations, see Methods.

### Comparison of Maketube genomes to standard Mtb genomes

We evolved a total of three times 30 genomes, corresponding to 3 different structural evolutions (3 populations of 10 individuals), from 3 different references: H37Rv (from L4 lineage), Beijing 18b (L2) and bovis AF2122/97 (animal lineage, related to L5], and this was done both with Maketube and with SNP-mutator (Set B, described above). To assess whether the parameters implemented in Maketube resulted in a sequence diversity comparable to that of natural strains, we aligned genomes from sets A (natural strains) and B to their reference using nucmer from the MUMmer tool [35, 46]. The same was done with SNP-Mutator genomes, with the expectation of no diversity and null distance to the reference. We computed the percentage of each genome not aligning to its reference and the reverse as proxies of their distance to reference. For natural genomes, when available, we chose the closest reference to the sample: 1) H37Rv for L4 strains, 2) 18b for L2 strains, and 3) AF2122/97 for *M. bovis* but also for L5 strains as L5 is related to the animal lineage. For L1 strains, we picked one reference (*M. bovis*), to represent the case of analysing strains from a lineage with no reference genome.

While SNP-Mutator genomes exhibited a distance to their reference of zero as expected, metrics derived from Maketube-evolved genomes fell globally in the range of natural genomes (median average distance to the reference of 0.62% and 0.51%, **Fig. 3A**). We also plotted each distance on separate axes. X-axis shows the combined impact of deletions in the sample, as they increase the relative length of the reference not aligning to the sample. Y-axis shows the impact of deletions in the reference (or their surrogate *i.e.* insertions of ancestral-like sequences in the sample), as they increase the length of the sample not aligning to the reference or the total length. Natural genomes showed a wide diversity in the proportion of deletions in the sample versus deletions in the reference, as expected due to the stochasticity of these rare events. Few outliers fitted expectations: BCG sample showed a higher distance of reference to sample (0.98%, X-axis) as expected due to the higher prevalence of large deletions in BCG lineage than in other *M. bovis,* and L5 sample showed a high distance to the affected reference, *M. bovis* (1.67%, Y-axis), as expected due to the higher frequency of deletions in *M. bovis* lineage as compared to L5 [15]. Maketube genomes exhibited more deletions in the reference (mean distance of sample to reference, 0.83%) than deletions in the sample (mean distance of reference to sample, 0.42%), due to parameters slightly favouring insertions of ancestral-like sequences as compared to deletions, and due to the marginal effect of the large duplication (∼0.03%), decreasing X-axis values due to the increase in length of the sample genome. Yet, the distance between Maketube genomes and their reference proved much more similar to that of natural genomes to a reference from the same or closest lineage, than what occurs for SNP-Mutator-like *in silico* evolved genomes: Maketube genomes, carrying structural variants as compared to their reference, appear much more realistic than SNP-Mutator-like genomes (**Fig. 3B**).

**Figure 3.**
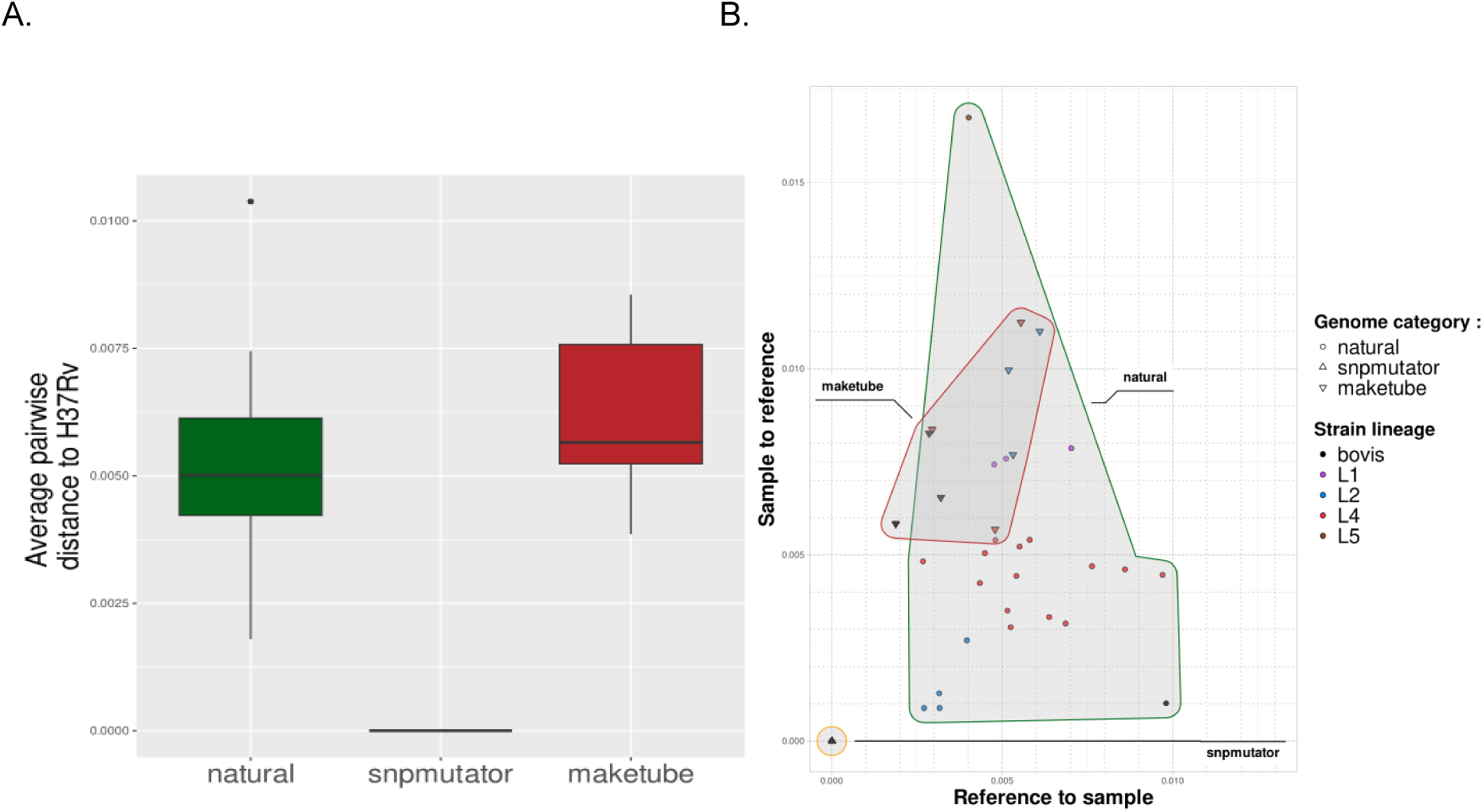
Distance between natural or artificial genomes and the closest reference genome. **A.** Mean of pairwise reciprocal distance between genomes and their reference as a function of their origin: “natural” refers to the genomes of natural strains reconstructed from a combination of long and short reads or derived from Sanger sequencing, “snpmutator” refers to SNP-Mutator-evolved genomes, and “maketube” refers to Maketube-evolved genomes. Distances were computed using genome length and aligned sequences (=similar sequences) output from the dnadiff wrapper based on the MUMmer nucmer tool (36). **B.** Plot of both distances, from sample to reference (X-axis) and reference to sample (Y-axis). A higher distance of sample to reference (Y-axis) is indicative of sequences in the sample not aligning to any region in the reference, *i.e.* regions deleted in the reference. A higher distance of reference to sample (X-axis) is indicative of sequences deleted in the sample genome. Natural genomes are circled in green and their lineage is colour-coded. *In silico-*evolved genomes derived from SNP-Mutator and Maketube are respectively circled in orange and red. The reference from which they were evolved, H37Rv, Beijing 18b, *M. bovis* is colour coded (respectively red, blue and black, as per the lineage they belong, *i.e.* L4, L2 and *M. bovis*). For Maketube genomes, as 3 different combinations of structural variants (*i.e.* 3 populations) were evolved for each reference, genomes from the same population lie at the same position so that only three different data points exist for each reference.

### Evaluation of genome aligners with Maketube genomes

Next, we wanted to make use of Maketube genomes to assess whether genome aligners, necessary when using natural *de novo* assembled genomes for benchmarking, might introduce biases in downstream analyses. As it is possible to directly backtrack the position of variants on the source genome, Maketube genomes are ideal to evaluate noise introduced by genome aligners. Using MUMmer nucmer and Minimap2, respectively used by Bush *et al*. and Marin *et al*. [6, 30], we aligned the 30 Maketube genomes evolved from H37Rv (Set B) to H37Rv, called variants, and compared the set of retrieved variants to the set of variants introduced by Maketube (**Fig. 1**). We proceeded similarly with SNP-Mutator genomes as the control.

As expected, on SNP-Mutator genomes, minimap2 and nucmer accurately found every SNP without identifying any false positives: precision (which measures here the ratio of true SNPs among all identified SNPs, *i.e.* is the complement of the false positive rate), and recall (which measures the ratio of true identified SNPs among all real SNPs, *i.e.* is the complement of the false negative rate) are both equal to one. Of note, this was true without excluding any part of the genome, keeping all repetitive sequences (**Fig. 4**). With Maketube genomes, for nucmer, the precision dropped only slightly but significantly below values obtained with SNP-Mutator genomes (mean of 0.9853 *vs* 1) while minimap2 retained a median precision of 1 (mean of 0.9999, Wilcoxon W = 0, p = 2,258e-12). When removing regions duplicated in the evolved genomes (that are single copy in the reference genome), the mean nucmer’s precision remained the same (0.9853), significantly lower than that of SNP-Mutator genomes (Wilcoxon W = 870, p= 2.26e-12, **S2 Table**, **Supplementary** Fig. 4). The largest parameter distinguishing Maketube and SNP-Mutator genomes was however not the precision but the recall of genome aligners. Mean recall for Minimap2 and nucmer were 0.9232 and 0.9549 respectively. This result was in part driven by duplicated regions. Without these, the mean recall of minimap2 was 0.9568 and the recall of nucmer of 0.9897. Nevertheless, these recall values, obtained with Maketube genomes without duplicated regions, are largely below 1 and significantly lower than those of SNP-Mutator genomes (nucmer, W = 855.5, p=8.341e-12; minimap2, W = 900, p=1.212e-12).

**Figure 4.**
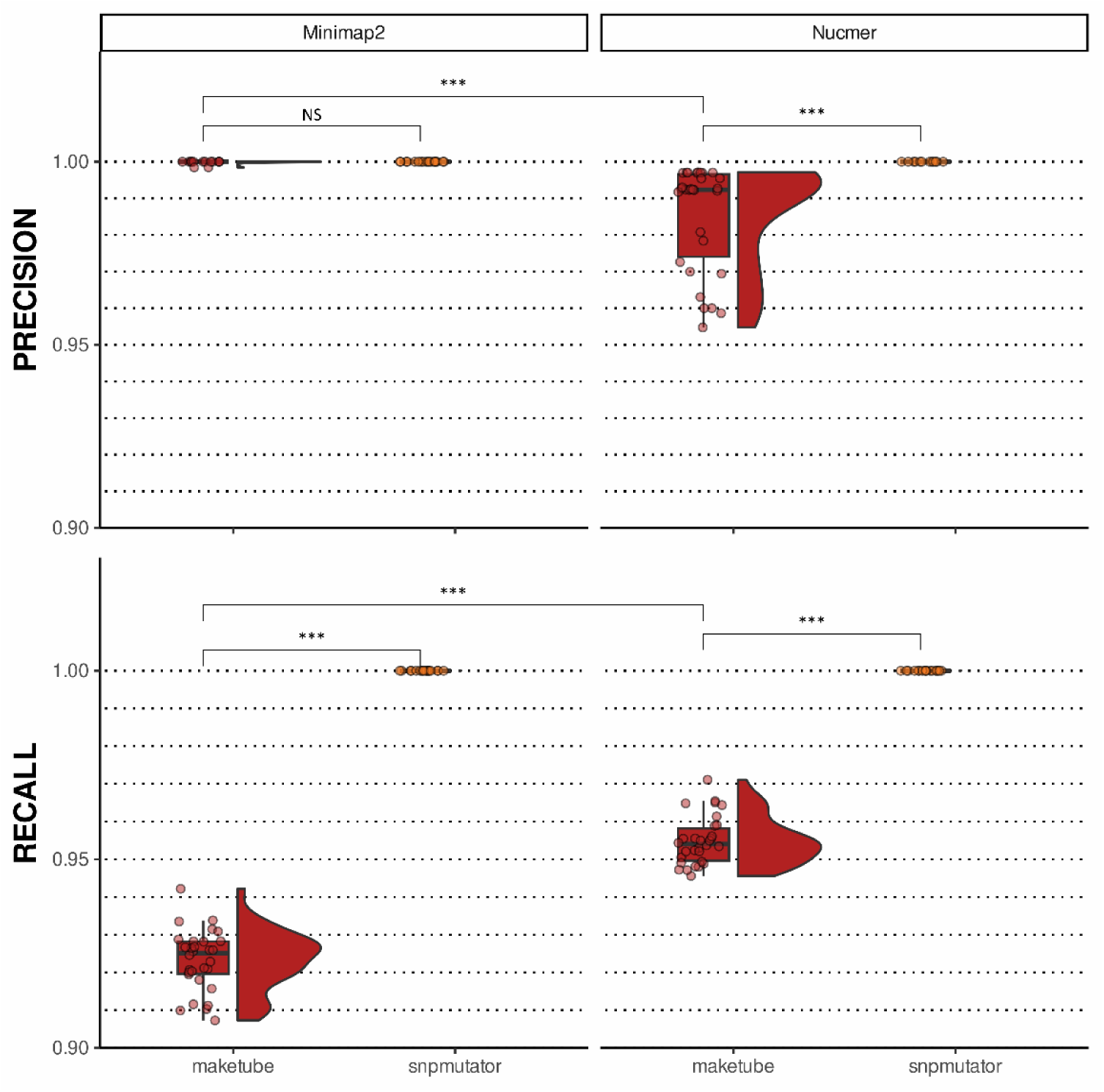
Precision and recall of genome aligners assessed using Maketube genomes. minimap2 PAF tool (left panel) and MUMmer3 nucmer (right panel) were tested for SNP recovery on Maketube-evolved and SNP-Mutator-evolved genomes. SNP-Mutator genomes represent positive controls for which SNP recovery is expected to be perfect. The level of statistical significance between variant calling strategies, taking into account Bonferroni corrections, is indicated by asterisks. *p<3.33×10⁻³; **p<3.33×10⁻⁴; *** p<3,33×10⁻⁵.

These results indicate that both nucmer and minimap2 do not recover all features of processed genomes, affecting mostly their recall: they both miss a subset of variants in all genomes. This will bias the downstream analyses in which they intervene.

### Benchmarking of three variant calling strategies on short reads using Maketube *in silico* evolved genomes

Next, we used Maketube-evolved genomes to benchmark three different variant calling pipelines and the corresponding variant callings without filtering, MTBseq *vs* samtools (raw), TBProfiler, and our own pipeline Genotube *vs* Freebayes (raw) (see Methods). We compared the metrics obtained to those derived from standard *in silico-*evolved genomes (SNP-Mutator, to the left, **Fig. 5**). To do so, we again used the 30 Set B genomes evolved from H37Rv, then we generated reads using Illumina ART and we aligned reads on H37Rv, taken as the reference genome (**Fig. 1**).

**Figure 5.**
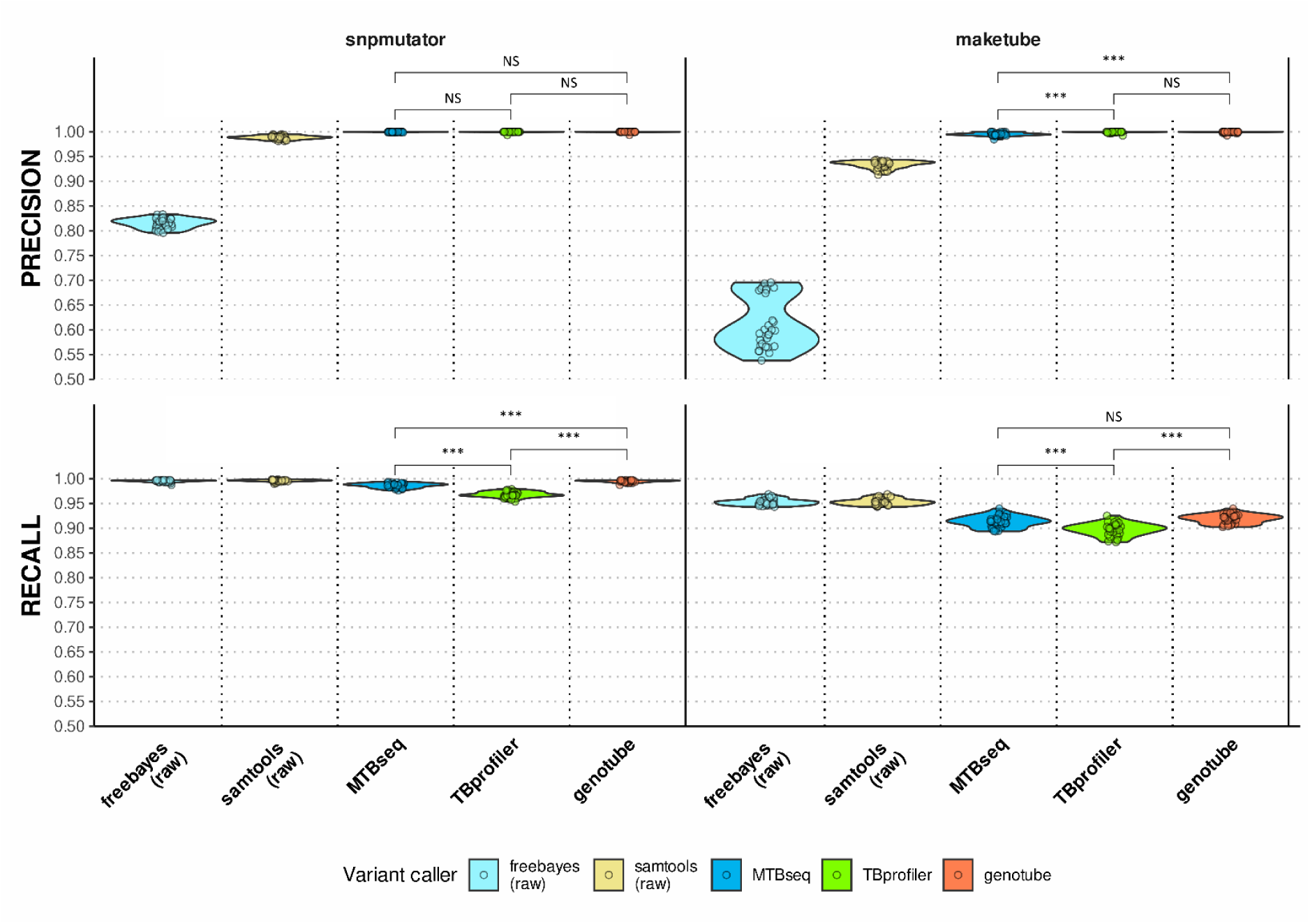
Precision and recall of the most popular variant calling strategies using SNP-Mutator and Maketube genomes with standard sequencing depth (30x). MTBseq, TBprofiler, and Genotube were assessed for recovering SNPs from reads aligned to H37Rv. We included performance on raw calls to highlight the importance of filtering processes (Freebayes raw calls for TBprofiler and genotube, and samtools raw calls for MTBseq). Statistically significant differences between variant calling strategies, taking into account Bonferroni corrections, are indicated by asterisks. *p<2,08×10⁻³ ; **p<2,08×10⁻⁴; *** p<2,08×10⁻⁵.

Performances of the three pipelines confirmed that filtering processes largely contribute to their precision, with moderate costs on the recall: precision was improved by almost 20% for Freebayes on SNP-Mutator genomes, and ∼35% on Maketube genomes (top of **Fig. 5**). When applied on SNP-Mutator genomes, all pipelines exhibited very high performances with mean precisions equal to 1 and very high recalls (respective mean recalls: MTBseq, 0.9865; TBprofiler, 0.9683; Genotube, 0.9948). In contrast, Maketube genomes discriminated pipelines’ ability to avoid any false positives, distinguishing TBprofiler and Genotube (mean precision of 1) from MTBseq (mean precision of 0.9954, MTB seq vs TBprofiler: W = 165, p = 2.60×10^-5^). MTBseq precision roughly corresponds to 0.5% false positives, *i.e.*, here, an average of 3 false positives for the average of 600 SNPs implemented in Set B. The gap in performance values on Maketube genomes *versus* SNP-Mutator was more important for the recall. TBprofiler had an average of ∼10% of false negatives instead of ∼3% (mean TBprofiler’s recall=0.8975 with Maketube genomes instead of 0.9675 with SNP-Mutator genomes). Similarly, other tools had a relatively low recall with Maketube genomes, exhibiting on average 9% of false negatives (MTBseq recall, 0.9133; Genotube recall, 0.9191, **Fig. 5**). One source of false negatives was the duplication region introduced in Maketube genomes. Without this region, the recalls of all 3 pipelines increased by ∼0.03 but stayed much below values of SNP-Mutator genomes (respective mean recalls without the duplicated region: TBprofiler, 0.9262; MTBseq, 0.9426; genotube, 0.9486).

### Impact of structural variations on variant calling using Maketube genomes

The specificity of Maketube genomes is to present all types of structural variants in contrast to standard *in silico*-derived genomes. We documented above how duplications affected the recall over the whole genome, but these could not explain all the performance drops observed between SNP-Mutator and Maketube genomes. We thus wanted to take a closer look at regions possibly responsible for additional loss of performance. We hypothesized that regions neighbouring structural variants could suffer from lower performance. During structural evolution, Maketube annotates regions within 300-bp (upstream and downstream) of structural variants, which facilitates such exploration. Regions surrounding the excision of an IS or a DR are labelled “IS scar regions” and “Deletion scar regions”. Regions surrounding ancestral-like insertions (those mimicking deletions in the reference) or the insertion site of an IS are respectively labelled “ancestral-like flanking regions” and “IS flanking regions” (**Fig. 2**). Regions without any structural variants are coined as “Neutral”.

Because these regions are much smaller than the complete genome, to gain sufficient power in our analyses, we generated set C consisting of 200 Maketube genomes evolved from H37Rv with a higher number of SNPs and indels (∼2,500 SNPs). We again generated reads using Illumina ART and aligned them to H37Rv. We called and filtered the variants using the Genotube methodology (**Fig. 1**). We compared the performance of variant calling in all region types listed above. As the size of the region has a large impact on precision and recall distributions (shorter regions having a much broader distribution simply due to the higher impact of stochasticity on small sample sizes, see for instance [47]), we subsampled 10 kbp of every region, which corresponds to an average of 6 SNPs in each of the 200 samples (see Methods for details).

In accordance with the high precision of the Genotube methodology described above, no significant precision loss was identified in the vicinity of any structural variants (**Fig. 6**). In contrast, recall on these small subsampled regions was lower in at least a subset of samples, materialized by a trail of rare values between 0.9 and 0.5, even in Neutral regions, but with largely different distributions. Recall close to 0.5 is due to rare false negatives among a low number of true positives. No statistically significant difference with Neutral regions was observed for deletion scar regions and IS scar regions, *i.e.* within regions flanking sequence loss in the focal samples. In contrast, sequence flanking DNA gains as compared to the reference, such as ancestral-like insertions and IS, significantly lowered the mean recall of their flanking regions compared to neutral regions (mean false negative rate of 9.8% in ancestral-like and IS flanking regions, *versus* 1.7% in Neutral regions).

**Figure 6.**
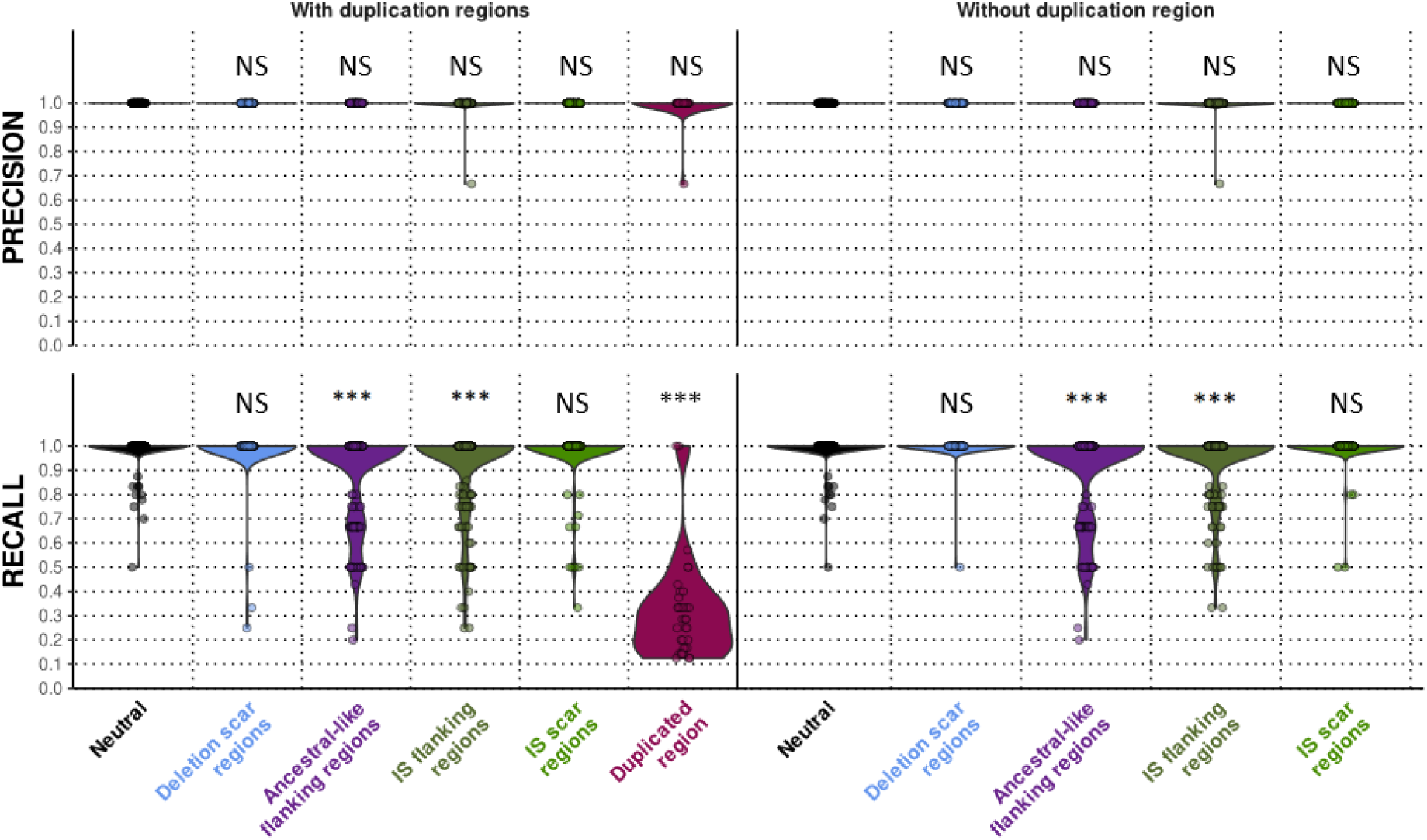
Precision and recall of variant calling nearby structural variants, assessed using Maketube genomes from set C (n=200). Precision and recall using the Genotube pipeline on different parts of the genome were explored as a function of the vicinity to structural variations (within 300 bp). Of note, as the variances in precision and recall are highly sensitive to the absolute number of variants in such short regions, we performed subsampling of each class to reach 10kb length for all. Statistically significant differences between specific regions and Neutral regions, taking into account Bonferroni corrections for significance, are indicated by asterisks. *p<3.125×10⁻³; **p<3,125×10⁻⁴; *** p<3,125×10⁻⁵.

## Discussion

We described here the Maketube pipeline, designed to build realistic Mtbc genomes evolved *in silico* from a genome of interest. The purpose was to have 100% reliable true variants to evaluate bioinformatics tools for genomics, such as variant calling pipelines. We built a total of 380 genomes evolved from 3 different source genomes, including H37Rv, forming 29 populations characterized by different sets of structural variants, and made them available on our GitHub. After confirming that they mimic relatively well real genomes, we used them to dissect the methods benchmarking variant-calling through reference based-alignment. We showed that benchmarking using *de novo* reconstructed genomes as templates are biased because they require genome aligners that provide non-exhaustive lists of true variants. We then showed that benchmarking based on standard *in silico* evolved genomes with no structural variants overestimate pipelines’ performance, especially their recall. Last, we confirmed the role of duplications in reducing pipelines’ performance and detailed the impact of other structural variants. In particular, flanking regions of sequences absent in the reference, due to insertions in the evolutionary branch of the sample such as IS insertions, or to deletions along the evolutionary branch of the reference genome, were found impacting variant-calling recall but not the flanking regions of sequences present in the reference and absent in the sample.

We discuss below: the realism of Maketube genomes and potential further applications, the properties of genome aligners, which outlines biases in benchmarking using *de novo* reconstructed genomes, some highlights deriving from our benchmark of MTBseq, TBprofiler and Genotube, the importance of our results for practice in variant calling in the clinics, and we provide some recommendations for variant-calling and for benchmarking of genomic tools, in particular the use of references other than H37Rv during reference-based variant-calling.

### Realism of Maketube genomes: limits and reliable applications

Maketube genomes are designed to evolve with events that closely resemble those affecting Mtbc genomes, with parameters that can be tuned for specific purposes. In the set B, parameters mimic the intra-lineage diversity: for instance, in genomes evolved from H37Rv, both the pairwise distance and the genome similarities were equivalent to those observed for L4 strains from different sublineages. In contrast, in set C, pairwise distances were inflated to gain power in detecting the variant calling performance in each type of genome regions, making them less realistic but more relevant to specific tests.

Behind the similarities with real genomes, even in set B, released Maketube genomes carry features that differ slightly from natural genomes, which were beneficial to our analyses but could affect other future analyses. First, we implemented a long duplication (150 kbp) in all our evolved genomes. The role of duplications is well acknowledged in Mtbc evolution, for instance for the PE-PPE gene family evolution [48]. Regarding recent evolution, very long duplications have been described, the longest being of 352,288bp in M41 strain [41]. Other cases of duplication concern a Beijing sublineage as described by Domenech et al., the duplication measuring around 350kb, BCG carrying a ∼100kbp duplication, the 60x amplification in tandem of a 2,440-bp region including *esxR/esxS* and flanking pe/ppe genes in a strain mutated in a repressor protein and evolved *in vitro* by Wang et al., the total amplification reaching 146.4kbp, shorter duplications including several direct variable repeats in the CRISPR locus, totalling up to 1-kb [45, 49–51]. Recently, among 16 Pacbio-sequenced genomes, one duplication of 7,3-kb was also identified [52]. Hence, even if duplications seem not pervasive, the focus on short reads and reference-based alignment made them overlooked in the last past years. Our choice of including one duplication in each evolved genome is in line with a better exploration of their potential role. When more data on frequency of duplications will be available, parameters could be adjusted to mimic them more realistically in Maketube. Second, we implemented only IS*6110* movements. Another insertion sequence exists and undergoes rare movements, IS*1081*. Movements of this sequence could be implemented but the type of consequences will be exactly the same as that of IS*6110*, so it will not reveal new types of errors during benchmarking of genomic tools. IS losses and IS gains could also be modelised. Third, we introduced artificial “Ancestral-like” sequences. As a reminder, these insertions mimic the deletions occurring on the branch of the reference genome. The total length of these insertions, of 13,335 to 38,948 bp in genomes from Set B, is in the range of deletions’ length, but slightly longer (10,570 and 15,634 bp in the same set). Of note, mainly due to the implemented duplication, released Maketube genomes are slightly longer than real genomes. Other small characteristics distinguishing Maketube genomes from real genomes include: Maketube genomes are handled as linear whereas real genomes are circular, the IS are perfectly cut and pasted, while natural transposition usually duplicates the CGA target sequence in which they insert [53]. For other developments, these features should be born in mind, or users may generate new Maketube genomes with parameters better fitting their needs. Improvements could also consist in implementing finer evolutionary processes, for instance to take genes into account in the evolutionary processes and to implement more realistic trees based on population dynamics such as progeny skew [54, 55].

As a summary, in their present form, released Maketube genomes are highly suited for benchmarking purposes such as variant calling of other short-read-based pipelines, of long-read sequencing including the detection of the exact position of structural variants, or for benchmarking of additional genome aligners. Any variant of interest can also be added manually to genomes carrying randomly picked variants, and thus its detection can be assessed in various contexts.

### Consequences of genome’ aligners errors on benchmarking using natural *de novo* reconstructed genomes

As explained previously, genome aligners such as MUMmer nucmer and minimap2 [6, 30] are used for benchmarking of variant-calling pipelines using *de novo* reconstructed genomes. Maketube genomes revealed that, while precision of nucmer and minimap2 was very good, recall of both tools was relatively disappointing: nucmer missed on average 5% of the positives against on average 8% for Minimap2 when dealing with genomes carrying a large duplication, and the regions with the highest recall with nucmer still missed more than 1.5% of the true variants. Benchmarking variant-calling tools with the resulting genomes, in which true SNPs have been missed, will impact what will be called true SNPs for the variant-caller: if they are also not detected by the variant-calling tool, these missed SNPs will mask false negatives of the variant-calling tool (false negatives will be underestimated); and if these SNPs are detected by the variant-calling tool, they will be spotted as false positives of the variant-calling tool (false positives will be overestimated). As a consequence, when comparing two or more variant-callers, the preference of one tool over the others might be simply due to its ability to reproduce genome aligner mistakes and not spot all real variants.

In addition, it must be remembered that benchmarking analyses based on natural *de novo* reconstructed genomes make the assumption that these natural genomes are perfectly reconstructed during the assembly, and that wet-lab experiments were free of mistakes and noise. Such events might add another layer of noise to the benchmarking results.

Altogether, these considerations underline the advantages of realistic *in silico-*evolved genomes such as Maketube genomes for benchmarking of genomic tools. If natural genomes are preferred, these analyses encourage a critical eye on the false positives and false negatives lists provided by different variant callers.

### Performance of assessed variant-calling tools

One objective of Maketube is to provide more reliable metrics when evaluating the bioinformatics pipelines used in the genomics of Mtbc and identify the most potent tools among their building blocks. We compared our in-house pipeline, Genotube, to two of the most commonly used TBprofiler, used both as a database and as an off-line pipeline, which is still updated in 2025, and MTBseq, published in 2018 and last updated in 2023. These pipelines implement different tools: whereas Genotube and the latest version of TBprofiler both use Freebayes and a similar approach to filtering (see material and methods), MTBseq implements the deprecated samtools’ mpileup, and GATK3 tools for base quality score recalibration and indels realignment [18, 19]. Analyses were performed on unmixed samples with reasonable depth of sequencing (30x on average), which corresponds to the lower end of coverage distribution of most recently sequenced samples [56].

Mainly thanks to SNP-filtering accompanying the variant calling process, almost all variants identified were genuine: Genotube and TBprofiler were highly precise, and MTBseq very seldom introduced a few false positives. This is in line with previous results preferring Freebayes over mpileup [6]. This very high precision was made at slight costs on the recall, with significant differences between the pipelines: even if the large majority of variants were called by all pipelines (∼95%), Genotube and MTBseq were more sensitive than TBprofiler, with TBprofiler pipeline losing up to 10% of the positives. This loss is triggered by structural variants, as seen by comparing analyses on genomes derived from Maketube and SNP-Mutator. Of note, the introduction of structural variants did not change the ranking of the three pipelines in either precision or recall.

Altogether, Genotube combines the highest precision of TB-Profiler and the better recall of MTBseq by using an intermediate level of filtering, together with the highly sensitive variant caller Freebayes. Genotube also includes a decontamination step which can be useful when dealing with contaminated or mixed samples. The filters implemented are compatible with lower depths than those applied in this study (variant detected on 5x coverage regions with 4 supporting reads, see Methods). Yet, Genotube does not implement any base-quality recalibration, potentially reducing its performance on contaminated samples, which should be independently benchmarked.

### A critical eye on past and future reference-based variant-calling inferences on clinical strains

Short-read WGS followed by reference-based variant calling is an increasing practice in clinical settings, due to the rapidity for obtaining relevant data for patient management. This process is eased by the development of all-in-one pipelines. The three main types of inferences are resistance prediction, lineage prediction, and clustering analyses [56]. We showed in this study that the claimed near-perfect precision of several of these all-in-one pipelines is confirmed with realistic *in silico-*evolved genomes *i.e.* there are almost no false positives. This means that output variants, for instance those associated with antibiotic resistance, are highly reliable. The only limit to the correct prediction of resistance when detecting a SNP is the knowledge on which SNPs actually cause resistance, a knowledge that is continuously increasing thanks to the wide efforts put on resistance catalogues. In contrast to the precision, we showed that the recall of all-in-one pipelines was previously overestimated: false negatives occur on average at a frequency close to 5%, but with a higher frequency near regions deleted in the reference genome and near insertion sequences. The following question is: should we reconsider previous analyses inferring absence of SNPs (*i.e.* sensitivity to antibiotics, when considering resistance prediction) because more than 1.5 to more than 8% (in case of 150 kbp duplication) of them may be false negatives? This could be the case if SNPs of interest lie in the vicinity of regions deleted in the reference genome and/or hot spots of IS insertions and/or lie in duplicated regions. We propose to explore these possibilities for several cases, always considering H37Rv as the reference genome.

First, regarding resistance, we may consider the case of Rifampicin (RIF) resistance as this antimycobacterial agent is the most potent anti-mycobacterial agent. This resistance is conveyed almost exclusively by short variants in an 81-bp region central to the *rpoB* gene and by nearby regions [57]. This RIF-Resistance Determining region (RRDR) is central to the 3,534 bp gene [58]. In addition, exhaustive studies on *rpoB* alleles show that the ancestral sequence of this gene is not a sequence longer than that of H37Rv [58]. This means that none of the most frequent variants conferring RIF resistance can be hidden by the nearby insertion of an IS, upstream or downstream the gene, or by a sequence present in the strain but absent in the H37Rv reference. Similarly, other short variants associated with drug resistance, such as isoniazid resistance due to *katG* mutation at position 315, fluoroquinolone resistance and ethambutol resistance, lie all at a distance over 300-bp from the gene borders as can be derived from their gene position and gene characteristics [57]. Concerning the insertion of sequences inside the genes, few unessential genes involved in resistance, such as *katG,* can carry viable IS insertions [59], but then it is the insertion itself that will trigger resistance, as it disrupts the protein involved in activating the prodrug, and not a potential variant hidden by the insertion. Last, we know of no case where drug-targeted genes are duplicated. Altogether, the finer knowledge we gained on the restricted limitations of classical variant calling tools confirms that drug resistance prediction derived from all-in-one pipelines is highly reliable, only limited by the extent of resistance catalogue and possibly by the poor attention put on rare IS insertions in non-essential genes.

Second, regarding the inference of clusters, all SNPs have to be considered to infer relatedness, and the standard practice considers a threshold of 5 SNPs in the distance between two samples to infer recent transmission [24, 60]. The revelation that recall is lower than previously advertised suggests that some strains inferred to be recently transmitted, might in fact be caused by more ancient transmissions. This might be particularly true for strains with a high number of IS (∼20 IS*6110*) such as many L4 and L2 strains. Still, the total sequence length in the vicinity of such insertions would be 300bp x2 x20 = 12 kbp. Compared to the global length of 4,411kbp, these sequences include only ∼0.3% of the potential SNPs. This means that inferences on recent transmissions for a small subset of strains are not strongly biased due to SNPs possibly hidden by insertions. Yet, the loss of recall would become detectable when applied to thousands of genomes. In addition, bias may further increase if samples carry large duplications. For finer interpretations, it is thus a recommendable practice to browse and inspect visually the borders of insertions to look for potential false negatives, and at SNPs important for clustering inferences. This also helps in dealing with other sources of variant-calling errors apart from bioinformatics ones, such as sample contamination, locally lower coverage, *etc*. [13, 14, 30].

The impact of repeated sequences such as subparts of PE-PPE gene families should theoretically have a stronger impact than insertions due to the large proportion (∼10%) of the genome they represent [61], both because they derive from duplications [62], and because these duplications cause insertions further impairing variant calling on their borders. These regions are commonly excluded during additional filtering processes [24], so if this practice is maintained, nothing is to be further explored. In contrast, if sequences among PE-PPE are kept to try and increase discriminatory power, finer exploration on alinements of reads on the genome is recommended as explained above.

Last, regarding lineage prediction, the first studies proposing classifications of SNPs included a single SNP for each lineage or sublineage [63]. If the SNP was missed, classification was, of course, missed. However, more recent classifications provide several SNPs per lineage and sublineage [16, 64, 65], which means that a small loss in recall will not impair classification.

Hence, we document here that the defects of variant calling outlined by Maketube genomes do not affect most clinically relevant features currently explored during WGS analysis of patient strains. But rising progress made on the detection of duplications and IS insertions, the better understanding of their impact on the physiology of the bacteria might change the landscape in the future, and variants neighbouring the structural variations may contribute to their impact and should not be overlooked.

### Pros and cons of adapting reference genome to the explored set

A recurrently outlined limit of reference-based variant calling is the use of a single reference. Indeed, only variants in sequences existing in the genome of reference can be detected. We show here that deletions in the evolutionary branch of the reference genome further impact variant calling, as well as the insertion of IS, by affecting the recall in the regions flanking the real or seeming insertion in the sample genome. What is the relative impact of these two origins of lower performance on variant calling in Mtbc, absence of regions and low recall on regions flanking insertions? Genome-genome alignments show that the average distance between two strains belonging to different lineages, such as a L2 strain and H37Rv reference lies below 1% (*i.e.* deletions total less than 1% of the total genome length). This means that analysing a L2 population of strains by H37Rv-based variant calling misses at most 1% of the variants. Second, regarding the impact of insertions, these will include IS movements, which are numerous in L2 strains. They will likely lead to 20 IS insertions. As each of these insertions leads to a total length of 600bp of flanking regions as defined in this study, the total length of sequences belonging to this class is 600 x 20 = 12,000 bp, *i.e.* 0.3% of the genome. Altogether, the strategy of working with the standard reference genome H37Rv on L2 strains should miss at most ∼1.3% of the variants as compared to working with a reference from the same lineage. As discussed above, these variants cannot be those triggering drug resistance. They could slightly impact inferences on clustering. Keeping H37Rv as a reference thus conveys a disadvantage regarding discriminatory power during clustering analyses, but this impact is low and compensated by safely detecting all variants affecting the phenotype, using the lists set up by the community. The choice of using a reference from the same lineage as the analysed samples, can be relevant if a high discriminatory power is searched for. As intralineage distance is around 0.5% on average, as exemplified on natural genomes in Fig. 3B, an average of 1 - 0.5 = 0.5 % more sequences will be scanned for variants. Regarding insertions, as IS are highly active in Mtbc, few IS remain at the same position even for strain belonging to the same lineage. This means that variants missed when using an L2 reference genome might miss the same range of variants due to IS as when using H37Rv. With the current knowledge, despite its seducing feature of taking into account the nature of the population of interest, changing reference for alignment-based variant calling to pick a reference from the same lineage seems of relatively poor interest. In contrast, the use of an ancestral sequence proposed by Harrison et al., with no deleted sequence, better fulfils this role [66]. An effort should be put to translate the resistance catalogue and all H37Rv annotations in this new genome.

### Beyond excluding regions: the structural variants that matter on variant-calling

Structural variants have already been pinpointed by other authors to affect variant-calling performance: Heupink defines “complex regions” as repeated regions including repeats, transposons, PE-PPE, phage genes, and duplicates, where variant calling is less efficient but to an extent that does not justify exclusion when handled with appropriate tools and filters [22]. In 2021, Bush et al. identified the presence of an indel as a risk factor for false positives. In 2024, Marin et al. documented that regions flanking structural variants exhibit substantial losses in precision and in recall, especially if these regions are in addition characterized by a low mappability. These structural variants were described as both insertions and deletions. In addition, Modlin et al showed that the GC content and sequencing biases may also impact variant-calling. Avoiding the negative impact of these regions is performed by excluding regions [6, 14, 22, 24, 30]. However, there is no consensual list on what sequences really need to be excluded.

Here, we showed that only duplicated regions and large insertions and not large deletions induce a significant drop in recall in a region the size of the read length around the point of insertion. Pipelines with lower precision and higher recall than those tested in this study could however behave slightly differently. These results are in line with the impact of structural variants identified by Bush and Marin, but point at a larger role of insertions as compared to deletions in inducing performance losses: we showed evidence that insertions in the sample genome as compared to reference-genome are the main driver of false negatives in variant calling. The impact of inserted sequences may in turn explain why different studies provided different lists of regions to be excluded: what is inserted in one genome as compared to the reference depends on its lineage and sublineage, and the exact position of all its IS. Hence, working on different samples may have contributed to different lists in regions to be excluded. This also means that any new study will come with its own region of lower reliability. Apart from regions that can be excluded like PE-PPE because of the large prevalence of duplications, it is safer to explore read alignments for any SNP critical for cluster inference or fine understanding of the genetic bases of phenotypes.

## Conclusion

To conclude, we developed a tool to build relatively realistic *in silico-*evolved Mtbc genomes (called Maketube genomes) and make available 380 such genomes for the community. These genomes are more suitable for performing genomic tools benchmarking than both simplistic genomes evolved with tools such as SNP-Mutator and natural genomes.

We showed that, for variant-calling benchmarking, the lower reliability and discriminatory power of SNP-Mutator genomes is due to their missing the impact of structural variants, and that the lower reliability of natural *de novo* reconstructed genomes is caused by the unavoidable use of error-prone genome aligners. This study thus further underlines the advantages of simulated genomes to capture real-life biases.

In addition, we provided new evidence that sequences absent in the reference impact variant detection in their flanking regions when using the current most popular tools for variant calling. Yet, in the absence of duplications, the effect size is relatively mild and should only be considered when trying to gain discriminatory power in molecular epidemiology studies or when exploring specific mechanisms involving IS movements. These small defects might, in addition, be reduced in the future by better variant calling tools. In turn, they could have a higher impact in the future if trying to exploit all sequencing information instead of excluding many regions due to lower reliability. In its course, this study led us to sum up evolutionary events affecting Mtbc genomes, which could be used for finer developments on simulated genomes in the future, incorporating more realistic dynamics of genome diversification.

## Methods

In short, Maketube genomes were first evolved structurally by introducing: 1) three deletions drawn from a gamma distribution (shape parameter=1, scale parameter=3,500 bp), based on the distribution of deletion regions sizes reported by Bespiatykh *et al*. [15]; 2) moving all 16 IS by a cut-paste process, 3) inserting ancestral-like regions reconstructed using random concatenation of kmers absent from H37Rv but derived from high quality genomes from set A using Jellyfish [67]), for a total length falling in the range 0.2 to 0.8% to mimic deletions in the reference with a total length targeted to be similar to total length of deletions in the sample but authorizing more events, 4) copying and pasting at the end of the genome a random sequence of 150kbp, in the range of duplications described by Weiner et al and half the size of longest described duplications [41]. After structural evolution, Maketube genomes were evolved introducing short variants using the Jackalope package from R [68], the GTR model derived from reliable genomes as per Guyeux et al, and listed in Suppl. Table 1 [69], an effective population size of 700 or of 2,500 depending on the total number of SNPs targeted, and a substitution rate of 1.23e-7 substitutions/site/year [70], in the range of mutation rates reported in various studies focusing on recent evolutionary rate. Indels were then introduced following a Lavalette distribution, calibrated from the same dataset, and with a ratio of 0.125 as compared to SNPs.

We generated “classical” *in silico* genomes using SNP-Mutator [26], with the same GTR model as that used in Maketube, and chose a total number of 800 SNPs and 100 indels.

Pairwise genome sequence comparisons presented in Fig. 3 were computed using the dnadiff wrapper of the nucmer from MUMmer3 [46]. Distances were computed as 1-A/L were A in the alignment of Genome 1 to Genome 2 output (standard alignment output from nucmer) and L the length of the Genome 2. The same version of the genome aligner was used for comparing the genome aligners’ performances (Fig. 4], to compare to Minimap2 paftools [35].

Reads were simulated from fasta genome sequences using ART with default parameters of MSv3 (MiSeq v3 built-in quality profile), a depth of 30X, a length of 300-bp for reads, and an insert length of 500-bp with 30-bp standard deviation [28]. Reads were aligned on the reference using bwa-mem2 [71], with a minimum mapping quality of 30. MTBseq (v1.1.0) was emulated, implementing the gatk3 indel realigner and base recalibrator, samtools mpileup to call the variants, and post-filtering kept only variants supported by at least 75% of reads and with at least 4 forward and 4 reverse reads [72, 73]. TB-Profiler (v6.6.3] was emulated by calling variant with Freebayes and post-filtering using bcftools view to keep only variants supported by at least 90% of reads, with at least 3 forward and 3 reverse reads [74]. Genotube, our in-house pipeline, was emulated by calling variants with Freebayes, and keeping only variants supported by 80% of the reads with a minimum of 5 reads, which implies a minimum of 4 reads supporting the SNP, and a quality > 30. Of note, before applying filters, single-nucleotide variants and indels were atomized and normalized using bcftools norm.

Statistical tests were performed in R. We used both the Wilcoxon signed rank test, reputed as more powerful, and the exact Welch permutation test, less sensitive to ties, with 100k permutations. Despite it being considered as too stringent, we considered Bonferroni’s correction by shifting the standard threshold of significance, to account for multiple testing [75]. Comparisons were considered significant only if the two tests detected a significant difference. When comparisons involved samples of very different sizes, such as for comparing the impact of structural variants, which would have conferred very different meanings for different comparison pairs, we applied subsampling.

Precision and recall were computed using the subsequent formula, where TP=True Positive, FP=False Positive, FN = False Negative ( Precision = TP/(TP+FP) ; Recall = TP/(TP+FN).

## Funding

ALM is supported by ANRS-ECTZ204875; this work was also supported by ANRS-ECTZ-190910 (GR and RZE).

## Author’s contribution

Conceptualization: RRDLV, ALM and GR; Software: ALM; Investigation: ALM; Analysis: ALM and GR; Writing: ALM, GR, RRDLV and RZE; Supervision: GR.

Competing interests: Authors declare to have no competing interests.

Data and materials availability: All data and codes are available on github.[75,76].

## Supplementary material

**Supplementary file 1 (S1). *Excel file.* Summary of strains info.** 1. List of all sample IDs from set A, 2. Statistics of genome characteristics from set B and set C strains, 3. Strains used to infer GTR model, 4. Exhaustive bioinformatic tools references.

**Supplementary file 2 (S2). *Excel file.* Results of statistical tests and exhaustive statistical data for** figure 4**-6.** 1. Output of all Wilcoxon and Welch permutation tests performed relative to Fig. 4 (comparison between nucmer and minimap2, for SNP-Mutator and Maketube genomes). 2. Output of all Wilcoxon and Welch permutation tests performed relative to Fig. 5 (comparison between pipelines, for SNP-Mutator and Maketube genomes). 3. Output of all Wilcoxon and Welch permutation tests performed relative to Fig. 6 (comparison between precision and/or recall of variant calling in different regions, for SNP-Mutator and Maketube genomes). 4. Absolute numbers of True Positives, False Negatives and False Positives relative to Fig. 6 (comparison between precision and/or recall of variant calling in different regions, for SNP-Mutator and Maketube genomes). 6. Additional tests performed when calling variants with TB-Profiler.

**Supplementary file 3. *Pdf.* Extensive methods (text).**

## Notes

https://github.com/adrienlemeur/maketube

https://github.com/adrienlemeur/maketube_supplemental

### Competing Interest Statement

The authors have declared no competing interest.

## References

1. Hunt M, Lima L, Anderson D, Bouras G, Hall M, Hawkey J, Schwengers O, Shen W, Lees JA, Iqbal Z. 2024. AllTheBacteria – all bacterial genomes assembled, available, and searchable. Bioinformatics, 10.1101/2024.03.08.584059.

2. Abdelwahab O, Torkamaneh D. 2025. Artificial intelligence in variant calling: a review. Frontiers in Bioinformatics 5.

3. Hall MB, Wick RR, Judd LM, Nguyen AN, Steinig EJ, Xie O, Davies M, Seemann T, Stinear TP, Coin L. 2024. Benchmarking reveals superiority of deep learning variant callers on bacterial nanopore sequence data. eLife 13:RP98300.

4. Derelle R, Madon K, Hellewell J, Rodriguez-Bouza V, Arinaminpathy N, Lalvani A, Croucher NJ, Harris SR, Lees JA, Chindelevitch L. 2025. Reference-Free Variant Calling with Local Graph Construction with ska lo (SKA). Molecular Biology and Evolution 42.

5. Koboldt DC. 2020. Best practices for variant calling in clinical sequencing. Genome Medicine 12.

6. Bush SJ, Foster D, Eyre DW, Clark EL, De Maio N, Shaw LP, Stoesser N, Peto TEA, Crook DW, Walker AS. 2020. Genomic diversity affects the accuracy of bacterial single-nucleotide polymorphism-calling pipelines. GigaScience 9.

7. Seah YM, Stewart MK, Hoogestraat D, Ryder M, Cookson BT, Salipante SJ, Hoffman NG. 2023. In Silico Evaluation of Variant Calling Methods for Bacterial Whole-Genome Sequencing Assays. Journal of Clinical Microbiology.

8. Olson ND, Lund SP, Colman RE, Foster JT, Sahl JW, Schupp JM, Keim P, Morrow JB, Salit ML, Zook JM. 2015. Best practices for evaluating single nucleotide variant calling methods for microbial genomics. Front Genet 6.

9. Riojas MA, McGough KJ, Rider-Riojas CJ, Rastogi N, Hazbón MH. 2018. Phylogenomic analysis of the species of the *Mycobacterium tuberculosis* complex demonstrates that *Mycobacterium africanum*, *Mycobacterium bovis*, *Mycobacterium caprae*, *Mycobacterium microti* and *Mycobacterium pinnipedii* are later heterotypic synonyms of *Mycobacterium tuberculosis*. International Journal of Systematic and Evolutionary Microbiology 68:324–332.

10. Senelle G, Sahal MR, La K, Billard-Pomares T, Marin J, Mougari F, Bridier-Nahmias A, Carbonnelle E, Cambau E, Refregier G, Guyeux C, Sola C. 2023. Towards the reconstruction of a global TB history using a new pipeline: TB-Annotator”. Tuberculosis 143.

11. Gagneux S. 2018. Ecology and evolution of *Mycobacterium tuberculosis*. Nature Reviews Microbiology 16.

12. Chiner-Oms A, Lopez MG, Moreno-Molina M, Furio V, Comas I. 2022. Gene evolutionary trajectories in *Mycobacterium tuberculosis* reveal temporal signs of selection. Proceedings of the National Academy of Sciences 119.

13. Goig GA, Blanco S, Garcia-Basteiro AL, Comas I. 2020. Contaminant DNA in bacterial sequencing experiments is a major source of false genetic variability. BMC Biology 18.

14. Modlin SJ, Robinhold C, Morrissey C, Mitchell SN, Ramirez-Busby SM, Shmaya T, Valafar F. 2021. Exact mapping of Illumina blind spots in the *Mycobacterium tuberculosis* genome reveals platform-wide and workflow-specific biases. Microbial Genomics 7.

15. Bespiatykh D, Bespyatykh J, Mokrousov I, Shitikov E. 2021. A Comprehensive Map of *Mycobacterium tuberculosis* Complex Regions of Difference. mSphere 6.

16. Napier G, Campino S, Merid Y, Abebe M, Woldeamanuel Y, Aseffa A, Hibberd ML, Phelan J, Clark TG. 2020. Robust barcoding and identification of *Mycobacterium tuberculosis* lineages for epidemiological and clinical studies. Genome Medicine 12.

17. Walter KS, Colijn C, Cohen T, Mathema B, Liu Q, Bowers J, Engelthaler DM, Narechania A, Lemmer D, Croda J, Andrews JR. 2020. Genomic variant-identification methods may alter *Mycobacterium tuberculosis* transmission inferences. Microbial Genomics 6.

18. Kohl TA, Utpatel C, Schleusener V, De Filippo MR, Beckert P, Cirillo DM, Niemann S. 2018. MTBseq: a comprehensive pipeline for whole genome sequence analysis of *Mycobacterium tuberculosis* complex isolates. PeerJ 6.

19. Phelan JE, O’Sullivan DM, Machado D, Ramos J, Oppong YEA, Campino S, O’Grady J, McNerney R, Hibberd ML, Viveiros M, Huggett JF, Clark TG. 2019. Integrating informatics tools and portable sequencing technology for rapid detection of resistance to anti-tuberculous drugs. Genome Medicine 11.

20. Jajou R, Kohl TA, Walker T, Norman A, Cirillo DM, Tagliani E, Niemann S, de Neeling A, Lillebaek T, Anthony RM. 2019. Towards standardisation: comparison of five whole genome sequencing (WGS) analysis pipelines for detection of epidemiologically linked tuberculosis cases. Eurosurveillance 24:1900130.

21. Feuerriegel S, Schleusener V, Beckert P, Kohl TA, Miotto P, Cirillo DM, Cabibbe AM, Niemann S, Fellenberg K. 2015. PhyResSE: a Web Tool Delineating *Mycobacterium tuberculosis* Antibiotic Resistance and Lineage from Whole-Genome Sequencing Data. Journal of Clinical Microbiology 53.

22. Heupink TH, Verboven L, Warren RM, Van Rie A. 2021. Comprehensive and accurate genetic variant identification from contaminated and low-coverage *Mycobacterium tuberculosis* whole genome sequencing data. Microbial Genomics 7.

23. Ezewudo M, Borens A, Chiner-Oms A, Miotto P, Chindelevitch L, Starks AM, Hanna D, Liwski R, Zignol M, Gilpin C, Niemann S, Kohl TA, Warren RM, Crook D, Gagneux S, Hoffner S, Rodrigues C, Comas I, Engelthaler DM, Alland D, Rigouts L, Lange C, Dheda K, Hasan R, McNerney R, Cirillo DM, Schito M, Rodwell TC, Posey J. 2018. Integrating standardized whole genome sequence analysis with a global *Mycobacterium tuberculosis* antibiotic resistance knowledgebase. Scientific Reports 8.

24. Meehan CJ, Goig GA, Kohl TA, Verboven L, Dippenaar A, Ezewudo M, Farhat MR, Guthrie JL, Laukens K, Miotto P, Ofori-Anyinam B, Dreyer V, Supply P, Suresh A, Utpatel C, van Soolingen D, Zhou Y, Ashton PM, Brites D, Cabibbe AM, de Jong BC, de Vos M, Menardo F, Gagneux S, Gao Q, Heupink TH, Liu Q, Loiseau C, Rigouts L, Rodwell TC, Tagliani E, Walker TM, Warren RM, Zhao Y, Zignol M, Schito M, Gardy J, Cirillo DM, Niemann S, Comas I, Van Rie A. 2019. Whole genome sequencing of *Mycobacterium tuberculosis*: current standards and open issues. Nature Reviews Microbiology 17.

25. Goossens SN, Heupink TH, De Vos E, Dippenaar A, De Vos M, Warren R, Van Rie A. 2022. Detection of minor variants in *Mycobacterium tuberculosis* whole genome sequencing data. Briefings in Bioinformatics 23.

26. Davis S, Pettengill JB, Luo Y, Payne J, Shpuntoff A, Rand H, Strain E. 2015. CFSAN SNP Pipeline: an automated method for constructing SNP matrices from next-generation sequence data. PeerJ Computer Science 1.

27. Kühl MA, Stich B, Ries DC. 2021. Mutation-Simulator: fine-grained simulation of random mutations in any genome. Bioinformatics 37.

28. Huang W, Li L, Myers JR, Marth GT. 2012. ART: a next-generation sequencing read simulator. Bioinformatics 28.

29 Li H. 2011. wgsim-Read simulator for next generation sequencing. Github repository.

30. Marin M, Vargas R, Harris M, Jeffrey B, Epperson LE, Durbin D, Strong M, Salfinger M, Iqbal Z, Akhundova I, Vashakidze S, Crudu V, Rosenthal A, Farhat MR. 2022. Benchmarking the empirical accuracy of short-read sequencing across theM. tuberculosisgenome. Bioinformatics 38.

31. Kolmogorov M, Yuan J, Lin Y, Pevzner PA. 2019. Assembly of long, error-prone reads using repeat graphs. Nature Biotechnology 37.

32. Hunt M, Silva ND, Otto TD, Parkhill J, Keane JA, Harris SR. 2015. Circlator: automated circularization of genome assemblies using long sequencing reads. Genome Biology 16.

33. Walker BJ, Abeel T, Shea T, Priest M, Abouelliel A, Sakthikumar S, Cuomo CA, Zeng Q, Wortman J, Young SK, Earl AM. 2014. Pilon: An Integrated Tool for Comprehensive Microbial Variant Detection and Genome Assembly Improvement. PLoS ONE 9.

34. Wagner J, Olson ND, Harris L, Khan Z, Farek J, Mahmoud M, Stankovic A, Kovacevic V, Yoo B, Miller N, Rosenfeld JA, Ni B, Zarate S, Kirsche M, Aganezov S, Schatz MC, Narzisi G, Byrska-Bishop M, Clarke W, Evani US, Markello C, Shafin K, Zhou X, Sidow A, Bansal V, Ebert P, Marschall T, Lansdorp P, Hanlon V, Mattsson C-A, Barrio AM, Fiddes IT, Xiao C, Fungtammasan A, Chin C-S, Wenger AM, Rowell WJ, Sedlazeck FJ, Carroll A, Salit M, Zook JM. 2022. Benchmarking challenging small variants with linked and long reads. Cell Genomics 2.

35. Li H. 2018. Minimap2: pairwise alignment for nucleotide sequences. Bioinformatics 34.

36. Khelik K, Lagesen K, Sandve GK, Rognes T, Nederbragt AJ. 2017. NucDiff: in-depth characterization and annotation of differences between two sets of DNA sequences. BMC Bioinformatics 18:338.

37. Dalquen DA, Anisimova M, Gonnet GH, Dessimoz C. 2012. ALF: A Simulation Framework for Genome Evolution. Molecular Biology and Evolution 29.

38. Batut B, Parsons DP, Fischer S, Beslon G, Knibbe C. 2013. In silico experimental evolution: a tool to test evolutionary scenarios. BMC Bioinformatics 14.

39. Pracana R, Burns R, Hammond RL, Haller BC, Wurm Y. 2022. Individual-based Modeling of Genome Evolution in Haplodiploid Organisms. Genome Biology and Evolution 14.

40. Gonzalo-Asensio J, Perez I, Aguilo N, Uranga S, Pico A, Lampreave C, Cebollada A, Otal I, Samper S, Martin C. 2018. New insights into the transposition mechanisms of IS6110 and its dynamic distribution between *Mycobacterium tuberculosis* Complex lineages. PLOS Genetics 14.

41. Weiner B, Gomez J, Victor TC, Warren RM, Sloutsky A, Plikaytis BB, Posey JE, van Helden PD, Gey van Pittius NC, Koehrsen M, Sisk P, Stolte C, White J, Gagneux S, Birren B, Hung D, Murray M, Galagan J. 2012. Independent Large Scale Duplications in Multiple M. tuberculosis Lineages Overlapping the Same Genomic Region. PLoS ONE 7.

42. Wang W-F, Lu M-YJ, Cheng T-JR, Tang Y-C, Teng Y-C, Hwa T-Y, Chen Y-H, Li M-Y, Wu M-H, Chuang P-C, Jou R, Wong C-H, Li W-H. 2020. Genomic Analysis of *Mycobacterium tuberculosis* Isolates and Construction of a Beijing Lineage Reference Genome. Genome Biology and Evolution 12.

43. Sanoussi CN, Coscolla M, Ofori-Anyinam B, Otchere ID, Antonio M, Niemann S, Parkhill J, Harris S, Yeboah-Manu D, Gagneux S, Rigouts L, Affolabi D, de Jong BC, Meehan CJ. 2021. *Mycobacterium tuberculosis* complex lineage 5 exhibits high levels of within-lineage genomic diversity and differing gene content compared to the type strain H37Rv. Microbial Genomics 7.

44. Brosch R, Gordon SV, Marmiesse M, Brodin P, Buchrieser C, Eiglmeier K, Garnier T, Gutierrez C, Hewinson G, Kremer K. 2002. A new evolutionary scenario for the *Mycobacterium tuberculosis* complex. Proceedings of the national academy of Sciences 99:3684–3689.

45. Brosch R, Gordon SV, Buchrieser C, Pym AS, Garnier T, Cole ST. 2000. Comparative Genomics Uncovers Large Tandem Chromosomal Duplications in*Mycobacterium bovis*BCG Pasteur. Yeast 1.

46. Kurtz S, Phillippy A, Delcher AL, Smoot M, Shumway M, Antonescu C, Salzberg SL. 2004. Versatile and open software for comparing large genomes. Genome biology 5:R12.

47. Hanczar B, Hua J, Sima C, Weinstein J, Bittner M, Dougherty ER. 2010. Small-sample precision of ROC-related estimates. Bioinformatics 26.

48. Fishbein S, van Wyk N, Warren RM, Sampson SL. 2015. Phylogeny to function: PE/PPE protein evolution and impact on *Mycobacterium tuberculosis* pathogenicity. Molecular Microbiology 96.

49. Domenech P, Kolly GS, Leon-Solis L, Fallow A, Reed MB. 2010. Massive gene duplication event among clinical isolates of the *Mycobacterium tuberculosis* W/Beijing family. Journal of bacteriology 192:4562–4570.

50. Wang L, Asare E, Shetty AC, Sanchez-Tumbaco F, Edwards MR, Saranathan R, Weinrick B, Xu J, Chen B, Benard A, Dougan G, Leung DW, Amarasinghe GK, Chan J, Basler CF, Jacobs WR, Tufariello JM. 2022. Multiple genetic paths including massive gene amplification allow *Mycobacterium tuberculosis* to overcome loss of ESX-3 secretion system substrates. Proceedings of the National Academy of Sciences 119.

51. Refrégier G, Sola C, Guyeux C. 2020. Unexpected diversity of CRISPR unveils some evolutionary patterns of repeated sequences in *Mycobacterium tuberculosis*. BMC genomics 21:841.

52. Stritt C, Reitsma M, Marin AMG, Goig G, Dötsch A, Borrell S, Beisel C, Comas I, Brites D, Gagneux S. 2025. Gene conversion and duplication contribute to genetic variation in an outbreak of *Mycobacterium tuberculosis*. Microbial Genomics 11.

53. Thierry D, Brisson-Noël A, Vincent-Lévy-Frébault V, Nguyen S, Guesdon J, Gicquel B. 1990. Characterization of a *Mycobacterium tuberculosis* insertion sequence, IS6110, and its application in diagnosis. Journal of clinical microbiology 28:2668–2673.

54. Stritt C, Gagneux S. 2023. How do monomorphic bacteria evolve? The *Mycobacterium tuberculosis* complex and the awkward population genetics of extreme clonality. Peer Community Journal 3.

55. Saber H, Almalahi MA, Albala H, Aldwoah K, Alsulami A, Shah K, Moumen A. 2024. Investigating a nonlinear fractional evolution control model using W-piecewise hybrid derivatives: An application of a breast cancer model. Fractal and Fractional 8:735.

56. Le Meur A, Zein-Eddine R, Lamer O, Hak F, Senelle G, Vernadet J-P, O’Donnell S, de la Vega RR, Refrégier G. 2024. Tools for short variant calling and the way to deal with big datasets, p. 219–250. In Phylogenomics. Elsevier.

57. Ismail NA, Nathanson C-M, Korobitsyn A, Zignol M, Kasaeva T, Rodwell T, Miotto P, Köser C, Walker T, Chindelevitch L. 2023. Catalogue of mutations in Mycobacterium tuberculosis complex and their association with drug resistance.

58. Williams DL, Spring L, Collins L, Miller LP, Heifets LB, Gangadharam PRJ, Gillis TP. 1998. Contribution of rpoB Mutations to Development of Rifamycin Cross-Resistance in *Mycobacterium tuberculosis*. Antimicrobial Agents and Chemotherapy 42.

59. DeJesus MA, Gerrick ER, Xu W, Park SW, Long JE, Boutte CC, Rubin EJ, Schnappinger D, Ehrt S, Fortune SM, Sassetti CM, Ioerger TR. 2017. Comprehensive Essentiality Analysis of the *Mycobacterium tuberculosis* Genome via Saturating Transposon Mutagenesis. mBio 8.

60. Walker TM, Ip CL, Harrell RH, Evans JT, Kapatai G, Dedicoat MJ, Eyre DW, Wilson DJ, Hawkey PM, Crook DW, Parkhill J, Harris D, Walker AS, Bowden R, Monk P, Smith EG, Peto TE. 2013. Whole-genome sequencing to delineate *Mycobacterium tuberculosis* outbreaks: a retrospective observational study. The Lancet Infectious Diseases 13:137–146.

61. Cole ST, Brosch R, Parkhill J, Garnier T, Churcher C, Harris D, Gordon SV, Eiglmeier K, Gas S, Barry CE, Tekaia F, Badcock K, Basham D, Brown D, Chillingworth T, Connor R, Davies R, Devlin K, Feltwell T, Gentles S, Hamlin N, Holroyd S, Hornsby T, Jagels K, Krogh A, McLean J, Moule S, Murphy L, Oliver K, Osborne J, Quail MA, Rajandream M-A, Rogers J, Rutter S, Seeger K, Skelton J, Squares R, Squares S, Sulston JE, Taylor K, Whitehead S, Barrell BG. 1998. Deciphering the biology of *Mycobacterium tuberculosis* from the complete genome sequence. Nature 393.

62. Gómez-González PJ, Grabowska AD, Tientcheu LD, Tsolaki AG, Hibberd ML, Campino S, Phelan JE, Clark TG. 2023. Functional genetic variation in pe/ppe genes contributes to diversity in *Mycobacterium tuberculosis* lineages and potential interactions with the human host. Front Microbiol 14:1244319.

63. Coll F, McNerney R, Guerra-Assuncio JA, Glynn JR, Perdigao J, Viveiros M, Portugal I, Pain A, Martin N, Clark TG. 2014. A robust SNP barcode for typing *Mycobacterium tuberculosis* complex strains. Nature Communications 5.

64. Coscolla M, Gagneux S, Menardo F, Loiseau C, Ruiz-Rodriguez P, Borrell S, Otchere ID, Asante-Poku A, Asare P, Sánchez-Busó L, Gehre F, Sanoussi CN, Antonio M, Affolabi D, Fyfe J, Beckert P, Niemann S, Alabi AS, Grobusch MP, Kobbe R, Parkhill J, Beisel C, Fenner L, Böttger EC, Meehan CJ, Harris SR, De Jong BC, Yeboah-Manu D, Brites D. 2021. Phylogenomics of *Mycobacterium africanum* reveals a new lineage and a complex evolutionary history. Microbial Genomics 7.

65. Guyeux C, Senelle G, Le Meur A, Supply P, Gaudin C, Phelan JE, Clark TG, Rigouts L, de Jong B, Sola C, Refregier G. 2024. Newly Identified *Mycobacterium africanum* Lineage 10, Central Africa. Emerging Infectious Diseases 30.

66. Harrison LB, Kapur V, Behr MA. 2024. An imputed ancestral reference genome for the *Mycobacterium tuberculosis* complex better captures structural genomic diversity for reference-based alignment workflows. Microbial genomics 10:001165.

67. Marcais G, Kingsford C. 2012. Jellyfish: A fast k-mer counter. Tutorialis e Manuais 1:1038.

68. Nell LA. 2020. jackalope: A swift, versatile phylogenomic and high-throughput sequencing simulator.

69. Guyeux C, Sola C, Noûs C, Refrégier G. 2021. CRISPRbuilder-TB:“CRISPR-builder for tuberculosis”. Exhaustive reconstruction of the CRISPR locus in mycobacterium tuberculosis complex using SRA. PLoS computational biology 17:e1008500.

70. Godfroid M, Dagan T, Merker M, Kohl TA, Diel R, Maurer FP, Niemann S, Kupczok A. 2020. Insertion and deletion evolution reflects antibiotics selection pressure in a *Mycobacterium tuberculosis* outbreak. PLoS Pathogens 16:e1008357.

71. Vasimuddin M, Misra S, Li H, Aluru S. 2019. Efficient architecture-aware acceleration of BWA-MEM for multicore systems, p. 314–324. In . IEEE.

72. DePristo MA, Banks E, Poplin R, Garimella KV, Maguire JR, Hartl C, Philippakis AA, Del Angel G, Rivas MA, Hanna M. 2011. A framework for variation discovery and genotyping using next-generation DNA sequencing data. Nature genetics 43:491–498.

73. Li H, Handsaker B, Wysoker A, Fennell T, Ruan J, Homer N, Marth G, Abecasis G, Durbin R, 1000 Genome Project Data Processing Subgroup. 2009. The sequence alignment/map format and SAMtools. bioinformatics 25:2078–2079.

74. Garrison E, Marth G. 2012. Haplotype-based variant detection from short-read sequencing. arXiv preprint arXiv:12073907.

75. VanderWeele TJ, Mathur MB. 2019. Some desirable properties of the Bonferroni correction: is the Bonferroni correction really so bad? American journal of epidemiology 188:617–618.

76. Le Meur A, 2025. Software Maketube. https://github.com/adrienlemeur/maketube.

75. Le Meur A, 2025. Supplementary information on Maketube manuscript methods. https://github.com/adrienlemeur/maketube_supplemental.

