## Supplementary material for "Improving the benchmark of variant calling in clonal bacteria using more realistic *in silico* genomes, the case of *Mycobacterium tuberculosis*": S3

Suppl. File 3 Extensive METHODS. Maketube manuscript.

### Generating Evolved Genomes with Maketube: Principles and Steps

Maketube evolves genomes from a source genome. This genome will later be used as a reference for aligning reads and performing variant calling. It is thus both a reference because it is the ancestor from which Maketube genomes are evolved, but it is also the reference in variant calling steps. H37Rv was used by default, but a set of genomes was also evolved from Beijing 18b and another one from *M. bovis* AF2122-97. Maketube may be used to evolve any other genome.
Maketube first generates structural variants, and then integrates short variants (both Single Nucleotide Variants referred to here as Single Nucleotide Polymorphisms or SNPs, and short insertions and deletions in the population, referred to as indels).

### Generation of structural variants, first step of Maketube genomes evolution

Evolutionary large modifications of Mtbc genomes include large deletions, inserting sequences (IS) jumps, and duplications. Large deletions occur both on the evolutionary branches of the studied samples and on that of the reference, they can in practice look like deletions or insertions. Hence, we implemented four types of structural variations in Maketube: the three types of standard evolutionary modifications listed above, plus insertions. These variations occur on the source to provide in its linear form, together with the list of IS positions to be moved provided in a separate file. The insertions require the user to provide a pool of kmers for constructing ‘ancestral-like regions’ ideally taken from reads or read fragments of the focal species that do not align to the reference genome. The tool performs sequential inclusion of variations and keeps track of regions affected by the variations, labelling them as “not available” so that no additional modification can impact them. Flanking regions (300bp upstream and downstream of each structural variant) are also labelled as “not available” and annotated to later test whether they exhibit features different from other regions. Only IS regions are initially labelled as ‘not available’.

First, the tool introduces deletions: it removes an adjustable number of regions from the reference genome, *i.e.* it implements Regions of deletion. In this study, we implemented 3 deletions in each structurally evolved genome. The size of each of these deleted regions is drawn from a gamma distribution (shape parameter=1, scale parameter=3,500 bp), based on the distribution of deletion region sizes reported by Bespiatykh *et al*. (2021). The location of deletions regions is chosen randomly across the genome. The flanking regions are annotated as “DR scars”.
Second, the tool performs the excision and the insertion at a different location of all IS*6110* insertion sequences. Although insertion sequences other than IS*6110* are described, IS*6110* is the one with the highest copy number and the most active. The initial positions of the inserting sequences are provided in a BED file. For this study, we identified the positions of IS*6110* in *H37Rv* using BLAST. Each IS*6110* is cut from the source sequence and then pasted in a random location within the genome. Their initial and end locations are stored in a partition file. The flanking regions are annotated as “IS scars” at the place of deletion, and “IS flanking regions” at the place of insertion.
Third, the tool inserts ancestral-like sequences (to mimic deletions occurring in reality on the evolutionary branch of the reference as explained above). Ancestral-like sequences are constructed by concatenating randomly selected kmers absent from the reference. Here, 31bp k-mers from all high-quality assemblies from set A (**Suppl Table 1**) were obtained using Jellyfish (Marçais et al., 2011). Kmers found in H37*Rv* were then removed. The targeted size

of total Ancestral-like sequences is selected in a range of 0.2 to 0.8% from the reference genome to mimic real data (Bespiyatykh et al 2021), with fragment sizes drawn from the same gamma distribution related to deletion characteristics (shape parameter=1, scale parameter=3500 bp). The flanking regions are annotated as “Ancestral-like flanking regions”. Fourth, the tool emulates duplications: Maketube copies and appends a single randomly-selected 150kbp sequence at the end of the linear version of the chromosome region. This size was chosen much below maximal duplicated regions' sizes (350 kbp, cf Weiner - 2012), but with a frequency higher than observed in natural genomes: we chose to include duplications in all evolved genomes, to have more power for detecting the consequences of such overlooked processes. The region flanking the duplication being single and short, we did not consider it for annotation and further analyses.

#### Generation of phylogenetic-based short variants, second step of Maketube genomes evolution

After incorporating structural variants, Maketube generates short variants. Several sets of short variants are incorporated in the same backbone that constitutes the structurally-evolved genome. We call “population” a set of genomes evolved from the same structurally evolved backbone. Incorporating these short variants first requires indicating a mutation rate, effective population size, and GTR model parameters as inputs. Using these parameters, and a genome, the **jackalope** R package (Nell, 2020) constructs a phylogenetic tree with branch lengths corresponding to the number of accumulated mutations. To estimate the parameters of the GTR model, we used a subsample of strains representative of the diversity from Guyeux et al. (2021) (**Supplementary Table 1**). We then counted the number of each mutation type and the number and size of small indels. Indels are introduced following a Lavalette distribution, calibrated to approximate the size of real indels, as per the described distribution of indels (Fletcher et al., 2009) from the same dataset. The number of indels was set as a ratio of SNPs, with a value of 0.125 according to their mean frequency in the genomes of reference, yielding one indel for every eight SNPs.

In a first set of Maketube genomes, belonging to set B (**Fig. 1B**), we aimed at mimicking the intra-lineage evolution which corresponds to a median accumulation of ~500 SNPs. We used the substitution rate of 1.23e-7 substitutions/site/year (Godfroid, 2020), in the range of mutation rate reported in studies focusing on recent evolutionary rate (Walker, 2013; Bryant, 2013; Roetzer, 2013), and an effective population size of 700. In our second set of genomes (**Fig. 1C**), we aimed at accumulating a sufficient number of variants to detect significant differences between regions. The effective population size was set to 2,500.

#### Backtrack and variant calling statistics

Maketube outputs fasta sequences and a VCF file with the variants of every haplotype. This VCF file lists all “true variants” corresponding to variants known to exist in the evolved genome. In addition, the script writes an evolution partition file that contains the list of every structural variant added to the source genome. Of note, since short variants are introduced into the evolved genome carrying sequences absent in the reference genome, some may not have an equivalent in the source genome. This occurs when an ancestral-like region or an inserting sequence is mutated. Such variants cannot be recovered through variant calling.

We developed a Python script called vcf2metrics to compare variants called after sequencing simulation and variant calling with the set of true variants. vcf2metrics identifies and sums true positives (TP), *i.e.* called variants that exist in the evolved genome, false positives (FP), *i.e.* called variants that are artifactual and do not exist in the evolved genome, and false negatives (FN), *i.e.* variants that exist in the evolved genome but were missed by the variant caller. It also computes precision (Precision= TP/(TP+FP) ) and recall ( Recall=TP/(TP+FN) ). Precision is an important metric because when inferring transmission, low precision indicates False Positives which leads to an overestimation of the pairwise distance between strains, *i.e.* underestimation of recent transmission. As a second metric, we chose to focus on recall (=TP/(TP+FN) ) that integrates FN instead of specificity that includes True Negatives (TN) (Specificity=TN/(TN+FN) ) for two reasons: first because when inferring transmission, if variants are not called (False Negatives), pairwise distances between strains will be underestimated and therefore recent transmission will be overestimated; second, False Negatives can lead to missing antibiotic resistance and not identifying to what lineage the strains belongs to (missing variant associated with resistance and missing variant used for lineage classification).

#### Structural variants terminology

We refer to the 300 bp downstream and the 300 bp upstream regions around a Deletion Region (outer borders) as the “DR scar regions”. The 300 bp downstream and the 300 bp upstream regions around the deletion left by the jump of Insertion Sequences are referred to as “IS scar regions”. The 600 (2x300) bp regions centred around the insertion site of Insertion Sequences are referred to as “IS flanking regions”. The 600 (2x300) bp regions centred around the insertion of Ancestral-like sequences are referred to as “ancestral-like flanking regions”. The number and size of structural variants are consigned in Supplementary File 1.

Because we introduced 3 deletions regions, the sum size of DR scar regions is 2 x 3 x 300 = 1.8kbp. As the number of ancestral-like regions is variable, the sum size of ancestral-like flanking regions is between 2 x 2 x 300 = 1.2kbp and 14 x 2 x 300 = 8.4 kbp. For H37Rv, we found 16 times the IS6110 region, so the sum size of IS flanking regions and IS scar regions are both 16 x 2 x 300 = 9.6 kbps.

#### “Classical” *in silico* genomes

We generated “classical” *in silico* genomes using SNP-Mutator. SNP-Mutator adds a desired number of short variants SNPs and indels of size 1 to a source genome, with characteristics derived from an evolutionary model provided by the user. We used the same GTR model (derived from natural genomes) as that used in Maketube. We ran SNP-Mutator 30 times with a fixed number of 800 SNPs and 100 indels on the three source genomes used to evolve Maketube genomes (**Fig1, B**). The tool outputs the list of true variants.

#### Comparing standard and Maketube-derived *in silico* genomes to natural genomes

Pairwise nucleotide sequence similarities to H37Rv reference genome were computed for artificial genomes (SNP-Mutator and Maketube-derived) and for natural strains genomes using

**dnadiff** from **MUMmer3** (Kurtz et al., 2004) (**Fig.1, A**). Two distances were computed. The first metric (Distance to the reference) is the ratio of the number of bases from the sample sequence that could not be aligned by nucmer on the reference sequence divided by the size of the reference sequence. The second metric 'Distance to sample' is the number of bases from the reference that could not be aligned by nucmer on the sample sequence divided by the size of the reference sequence.

As a consequence, a sequence present in the studied genome but absent in the reference (such as an ancestral-like region) will increase the first metric but not the second and conversely, a deletion in the studied genome (deletion region) will lower the second metric without affecting the first.

#### Assessing the performance of genome aligners

We aligned the genomes evolved *in silico* with maketube and SNP-Mutator on their reference using MUMmer3 and minimap2. For MUMmer3, we used the dnadiff wrapper of nucmer and the output was converted to VCF using all2vcf (Schiavinato et al., 2022). For minimap2, variants were called using paftools using its default parameters (**Fig. 1A2**). We then compared the recovered variants to the list of variants of the evolved genomes.

#### Comparing the performance of three variant calling strategies using short reads derived from *in silico* genomes

From the 30 SNP-Mutator and 40 maketube genome fasta, we generated FASTQ archives using **Illumina ART** (Huang et al., 2012) with default parameters of MSv3 (MiSeq v3 built-in quality profile), a depth of 30X, a length of 300bp for reads, and an insert length of 500bp with 30 bp standard deviation.

Reads were directly aligned to the reference genome using **bwa-mem2** (Vasimuddin et al., 2019), with a minimum mapping quality of 30. Of note, trimming was not applied because it makes no sense in this case as no adapters are added by ART. Trimming was in addition found to have a rare impact on variant calling on real data thanks to the improved versions of read alignment procedures (Bush et al, 2020).

Variants were then called independently with the following pipeline :

1. MTBseq (v1.1.0) was run with default parameters. This pipeline uses bwa mem to align the read, the gatk3 indel realigner and base recalibrator, and samtools mpileup to call the variants. Only variants supported by at least 75% of reads and with at least 4 forward and 4 reverse reads are retained. Because MTBseq does not output vcf but tables by default, the variant calling and filtering steps were emulated with the same tools in the same versions in order to get a VCF file.
2. TB-Profiler (v6.6.3) (Verboven et al., 2022) was run with default parameters but without trimming the reads. Reads are aligned with bwa mem, and variants called with freebayes. Because TB-profiler does not output VCF files, the filtering steps were replicated using bcftools view with the same filter thresholds (variant supported by at least 90% of reads, with at least 3 forward and 3 reverse reads) (Garrisson et al., 2024).
3. Genotube, our in-house pipeline, with default parameters, but without trimming the reads and decontamination preprocessing of the reads. Variants are called with

freebayes and only variants supported by 80% of the reads with a minimum of 5 reads and a quality > 30 were kept.

Variants and indels were atomized and normalized using **bcftools norm**.

**Statistical analyses**

In order to compare the performance of two methodologies, we performed two statistical tests:

- A two-sided Wilcoxon signed rank test, when the two distributions are not independent (such as the precision of two variant callers on the same set of strains). Under H0, the distribution of X and Y differ by a fixed location shift μ.

- a two-sided permutation test using the Welch statistic. Under H0, X and Y come from the same distribution.

The permutation test is more reliable than the Wilcoxon test when many values are equally ranked.

All tests were performed in R. We applied the Bonferroni correction for every experiment. Differences are described as significant when they affect both tests. The detail of every test performed for each experiment is given in Supplemental Table 2.

When comparisons involved samples of very different sizes such as for comparing the impact of structural variants, which would have conferred very different meanings for different comparison pairs, we applied subsampling.
